# Robust pan-junctional reinforcement preserves the gut epithelial barrier under mechanical stress

**DOI:** 10.1101/2025.10.04.680144

**Authors:** Vishnu Krishnakumar, Amalia Ferran, Sandra Bernat-Fabre, Justine Creff, Nicolas Cenac, Denis Krndija

## Abstract

Epithelia are specialized tissue barriers that safeguard the organism’s internal milieu from the hostile external environment, a function critically dependent on intercellular junctions. In the colon, this barrier is repeatedly challenged by mechanical distension from faeces and it is unknown whether and how the colon adapts to such stress, which could otherwise compromise barrier integrity. Here, we show that faeces-mediated mechanical distension triggers coordinated remodelling at cell- and tissue-scale, suggestive of mechanoadaptation. This response includes recruitment of junctional proteins at all three types of adhesive cell-cell junctions (tight junctions, adherens junctions and desmosomes). We identified two modes of recruitment: stable responders (ZO-1, E-cad, plakoglobin) with sustained junctional enrichment, and adaptive responders (desmoplakin, keratin 8) with progressive accumulation during distension. Distension was also accompanied by perijunctional recruitment and activation of non-muscle myosin II (NMMII). Through genetic, pharmacological, and mechanical perturbations, we demonstrated that NMMII activation is an early and critical step for mechanoadaptation. This process requires extracellular calcium influx, and Piezo1 activation is sufficient to trigger NMMII activation and junctional recruitment. Loss of NMMII function abrogated the junctional response to distension across all three junctional complexes, including desmosomes, resulting in disorganised junctions and barrier breach. Together, our findings uncover a robust physiological mechano-adaptive response of the adult colonic epithelium to an extrinsic mechanical stress, whereby coordinated reinforcement of all junctional complexes, controlled by myosin II and mechanosensitive calcium influx, plays an essential role in preserving intestinal barrier integrity.

## Introduction

Epithelia are the specialised tissue barriers that protect the internal environment of an organism from the hostile external environment, thereby ensuring normal tissue function and homeostasis. The gut epithelium must withstand multiple environmental insults, including diverse microbiota, toxins, and mechanical forces (e.g., compression and stretch) arising from distension by luminal contents and muscle-driven peristalsis (*1*–*3*). Despite these challenges, the gut epithelium remarkably maintains its critical roles in nutrient and water absorption while preserving barrier integrity throughout life. The adult colon exhibits a distinct architecture in which the mucosa – comprising the epithelium and underlying lamina propria – folds into the lumen and is organized into two main regions: crypts, which contain proliferative stem and progenitor cells, and plateaus, the intercryptal regions lined with differentiated epithelial cells directly exposed to luminal contents. However, how this mucosal architecture physically adapts to the extrinsic mechanical load imposed by faeces *in vivo* – and, in particular, how the colonic epithelium preserves barrier integrity to maintain homeostasis and prevent pathogen translocation – remains poorly understood. Epithelial tissue integrity is maintained by adhesive cell-cell contacts, comprising tight junctions (TJs), adherens junctions (AJs) and desmosomes. TJs form the most apical junctional complex, physically sealing the paracellular space through the transmembrane proteins occludin and claudins, and play key roles in regulating apico-basal polarity and selective ion permeability (*4*). They are mechanically linked to the underlying actomyosin belt via the Zonula Occludens (ZO) family of scaffolding proteins (*5*). In particular, ZO-1 responds to intrinsic mechanical forces arising during cellular processes through recruitment to the junctions (*6, 7*). AJs are located beneath the TJs and mechanically integrate epithelial cells through coupling of cadherins to perijunctional actomyosin belts via catenins and cytoplasmic linkers. Studies in simple epithelia and developmental models have shown that AJ proteins respond to mechanical tension by clustering, undergoing conformational changes, and engaging the contractile actomyosin network, thereby reinforcing junctional complexes to maintain tissue cohesion (*8*–*11*). Desmosomes, positioned basolaterally beneath AJs, provide mechanical stability by anchoring to intermediate filaments (*12*–*14*). In contrast to TJs and AJs, however, very little is known about whether desmosomes can sense or adapt to extrinsic forces *in vivo*, or how they might interact with the actomyosin cytoskeleton. This gap in knowledge is especially striking given their central role in mechanical resilience.

During morphogenesis, epithelial cells employ multiple strategies to sense mechanical forces, including adhesion complexes and mechanosensitive ion channels, which may trigger junctional remodelling to preserve tissue integrity (*6, 7*). Yet in adult epithelia *in vivo*, it remains poorly understood how extrinsic forces are sensed and how adhesive complexes—particularly desmosomes—and their coupling to actomyosin and intermediate filaments contribute to tissue integrity and barrier maintenance.

Here, we investigated how the colonic epithelium responds to physiological mechanical forces by combining high resolution 3D tissue imaging with mechanical and pharmacological perturbations in both *in vivo* (mouse models) and *ex vivo* (gut explant) systems. We found that TJs, AJs and desmosomes respond to extrinsic mechanical force through recruitment of their respective proteins, with distinct kinetics. Desmosomes, in particular, displayed progressive reinforcement during distension, highlighting their adaptive mechanoresponse in the gut. Non-muscle myosin II emerged as a critical mediator of this junctional recruitment under mechanical stress – its inhibition or depletion impaired force-induced recruitment across all the junctional complexes and led to barrier defects. Finally, we propose that physiological mechanical forces are sensed via mechanosensitive ion channels, including Piezo1, since modulation of extracellular calcium influx was sufficient to trigger or abolish this response and myosin II activation.

## Results

### 1. Faeces-mediated distension of colon leads to cell- and tissue-scale epithelial remodelling and junctional recruitment

To investigate the impact of mechanical forces in the adult murine colon, we first assessed the organ’s gross morphology with respect to the presence of faecal pellets. Regions containing faeces, interspersed with faeces-free regions, appeared visibly distended (**Fig. 1A**). By confocal imaging of colonic thick sections, either distended or non-distended by faeces, we observed unfolding of mucosal folds specifically in the distended regions (**Fig. 1B**). Distension was associated with tissue-level morphological changes in crypt and plateau organization: the inter-cryptal plateaus were expanded, whereas the crypts appeared shorter (**Fig. 1B, D**). Since the plateau regions are in direct contact with the gut lumen and faeces – and thus more exposed to luminal contents and microbiota – we focused our detailed morphometric analyses on these regions. Epithelial cell aspect ratio was altered (34% decrease) (**Fig. 1C, E**), suggesting that the colonic epithelial cells undergo significant deformation and strain upon faecal distension. To understand how the epithelial cells respond to the distension-induced strain, we assessed the adhesive cell-cell junctions (tight junction (TJ), adherens junction (AJ) and desmosomes) by immunofluorescence on thick sections with luminal side facing up (*en face*), enabling high-resolution 3D imaging of the cell-cell junctions. TJ proteins (ZO-1, occludin), along with AJ protein E-cadherin (E-cad) were enriched at the junctions in faeces-distended regions (2.4, 2.4 and 1.8-fold, respectively), compared to the adjacent, non-distended regions (**Fig. 1F, I, J; Fig. S1A, B**).

**Fig. 1.**
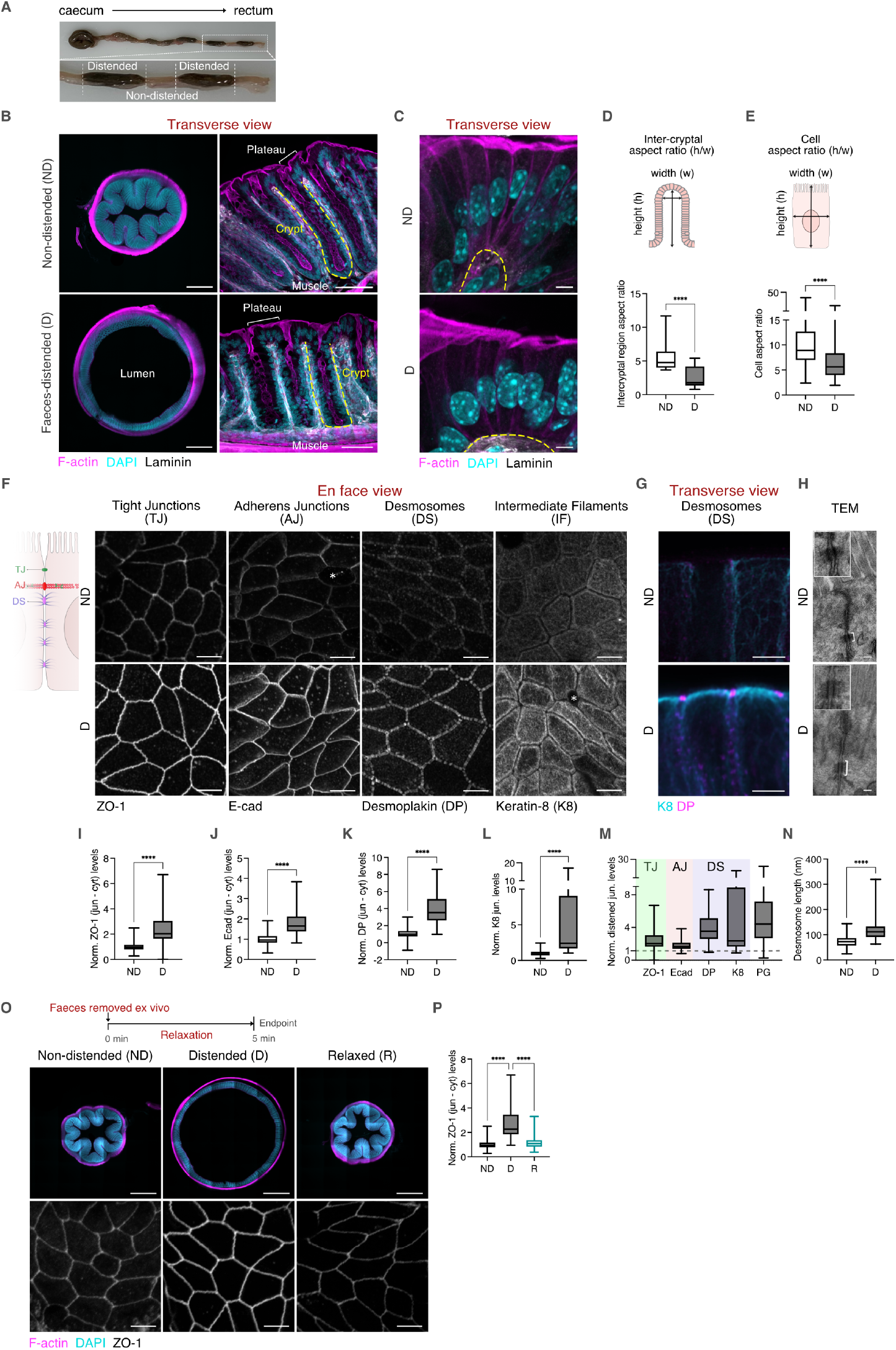
Faeces-mediated colonic distension leads to tissue remodelling and junctional recruitment. **(A)** Representative image of a mouse colon aligned from proximal (caecum) to distal (rectum) ends, showing non-distended regions (ND) without faeces and distended regions (D) with faeces. **(B)** Representative images of thick section of mouse colonic tissue in transverse view, showing lumen, plateau regions (white brackets), muscle layers and crypts (outlined in yellow dashed lines), and stained for F-actin (magenta), DAPI (cyan) and laminin (grey) in both ND and D regions. The left panel is a tiled confocal reconstruction. Scale bars: left panel, 500 µm; right panel, 50 µm. **(C)** Representative transverse images of plateau regions in the colonic epithelium, stained for F-actin (magenta), DAPI (cyan) and laminin (grey) in both ND and D region. Basal side of the cell outlined as yellow dashed line. Maximum Z-projections (2-4 µm range). Scale bar: 5 µm. **(D)** Top: Scheme showing parameters used for measuring the intercryptal aspect ratio. Bottom: Box plots showing intercryptal aspect ratio in ND and D plateau regions, (n= 5-9 intercryptal regions/condition, N=4 independent experiments) **(E)** Top: Scheme showing parameters used for measuring the cell aspect ratio. Bottom: Box plot showing cell aspect ratio in ND and D plateau regions, (n>28 cells/condition), N=4 independent experiments. **(F)** Left: Scheme showing adhesive cell-cell junctions in colonic epithelium. Right: Representative *en face* images of colonic epithelium, stained for tight junction (TJ) protein ZO-1, Adherens junction (AJ) protein E-cad, and desmosomal (DS) protein Desmoplakin (DP) and desmosome-associated intermediate filament (IF), K8 in both ND and D regions. Maximum Z-projections (1-3 µm range). Scale bar: 5 µm. **(G)** Representative transverse images of colonic epithelium, stained for K8 (cyan) and DP (magenta) in both ND and D regions. Maximum Z-projections (2-4 µm range). Scale bar: 5 µm. **(H)** Representative transverse TEM images showing desmosomes (electron-dense regions, white bracket or arrow; enlarged insets at top left corner) in ND and D regions. Scale bar: 100 nm. **(I-L)** Box plots showing junctional intensity for ZO-1 (L), E-cad (M), DP (N) and K8 (O) levels in ND and D regions. (n=34-85 junctions/condition, N=4-9 independent experiments). **(M)** Comparative box plot showing fold changes in junctional protein levels of ZO-1, E-cad, DP, K8 and PG in the D regions. Dotted line represents the normalised average for ND. **(N)** Box plot showing desmosome length in ND and D colonic epithelium (n=7-27 desmosomes/condition, N=3 independent experiments **(O)** Top panel: Scheme showing experimental approach for relaxation assay. Middle panel: Representative stitched transverse confocal images from tiled acquisitions of colonic epithelium stained for F-actin (magenta) and DAPI (cyan), showing the non-distended (ND), distended (D) and relaxed (R) regions. Scale bar: 500 µm. Bottom panel: Representative *en face* images of colonic epithelium stained for ZO-1, showing ND, D and R regions. Maximum Z-projections (1-3 µm range). Scale bar: 5 µm. **(P)** Box plot showing junctional ZO-1 intensity in ND, D and R regions. (n=60-80 junctions/condition, N=3 independent experiments). Statistical analyses were performed using the Mann-Whitney U test (D), (E), (I-L) and N or Kruskal-Wallis test followed by Dunn’s multiple comparison (P). Significance is denoted as: ****p < 0.0001, ***p < 0.0005, ** p < 0.005. ND: Non-distended, D: Distended.

Desmosome-associated proteins (desmocollin (DC), plakoglobin (PG), desmoplakin (DP) and keratin 8 (K8)) were also enriched at the junctions (**Fig. 1F, K, L; Fig. S1C, D**) with DP, PG and K8 showing even higher recruitment (3.9, 5.3, and 4.8-fold, respectively) upon distension when compared to the TJ and AJ proteins (**Fig. 1M**). The desmosomal adaptation (DP and K8) upon distension was also evident in transversal colonic sections as apical enrichment (**Fig. 1G**), as well through a significant increase in the length of desmosomal plaques via transmission electron microscopy (TEM; non-distended: 74 nm; distended: 117 nm; (**Fig. 1H, N**). In addition to the apical-most desmosomes analysed in our study, we also observed enrichment of desmocollin, desmoplakin, and keratin 8 in other desmosomes along the basolateral membrane in faeces-distended regions (**Fig. S1E, F**).

In stark contrast, in non-distended regions desmosomal proteins (PG and DP) were primarily cytoplasmic with only weak junctional staining, whereas in distended regions they exhibited striking cytoplasmic clearance and strong junctional enrichment, consistent with force-mediated recruitment from the cytoplasm to the junctions (**Fig. S1G-K**).

Importantly, we did not observe any difference in total protein levels of key junctional markers between faeces-distended and non-distended regions, suggesting a force-dependent relocalisation of junctional proteins rather than *de novo* synthesis (**Fig. S2A, B**). In contrast to most of the junctional proteins we studied, beta-catenin, an AJ-associated protein, did not show any junctional recruitment upon distension, pointing to a selective protein recruitment to the junctions upon force (**Fig. S2C**). The enrichment of junctional proteins at cell-cell contacts, together with comparable total protein levels in distended and non-distended regions, strongly suggests that this response arises from force-dependent redistribution of pre-existing proteins rather than de novo synthesis, as previously shown for E-cadherin, which can be delivered through vesicle trafficking or lateral diffusion and clustering within the membrane (*10, 15*–*17*). Interestingly, under basal conditions, plakoglobin and desmoplakin displayed diffuse cytoplasmic localisation; upon tissue distension, they were cleared from the cytoplasm and enriched at junctions, resembling the force-mediated relocalisation of ZO-1, previously attributed to phase separation under mechanical force (*6*).

To explore the stability of the accumulated junctional complexes and decipher whether the distension is required to maintain the recruited proteins at the junctions, we analysed gut fragments with/without faeces and upon faeces removal *ex vivo*. Upon faeces removal, we observed re-folding of mucosal folds and dissociation of proteins (ZO-1, PG and DP) from the junctions within 5 min (**Fig. 1O-P; Fig. S2D-I**). Taken together, these data suggest that the colon responds by adaptation to the extrinsic physiological mechanical force at multiple scales, as evidenced by the tissue and cell-level epithelial remodelling and junctional recruitment of adhesive TJ, AJ and desmosomal proteins.

### 2. Time-resolved colonic distension *in vivo* reveals differential junctional recruitment kinetics and transient apical constriction

The duration of faecal residence and transit through the mouse colon is not well characterised and may depend on multiple factors, including stress levels, diet and microbiota (*18, 19*). Additionally, faecal pellets are variable in size (*20*), which we also observed in our measurements of pellet size (**Fig. S2J, K**). To achieve controlled distension with precise temporal resolution and consistent volume, we introduced an inflatable catheter into the distal colon of anesthetised mice (*21*) (**Fig. 2A**). To investigate the recruitment kinetics of junctional complexes, colonic distension was performed at a constant inflation volume and maintained for 5, 15 or 30 min. These intervals were selected to cover short to more sustained stimulation, as the residence time of faecal pellet at a given site is difficult to determine directly. Estimates of mouse colonic transit are around 2 h (*22*) but this likely varies with faecal viscosity and consistency (*23*), suggesting that faecal distension can exert pressure on the colonic wall over extended periods. Catheter-mediated distension induced alterations in cell aspect ratio and intercryptal morphology that were comparable to those observed with faeces-mediated distension (**Fig. S3A, B**). Consistent with faecal distension, ZO-1, E-cad and PG showed significant junctional recruitment upon catheter-mediated distension, which was evident from 5 min (the earliest time point) and remained stable for up to 30 min (**Fig. 2B, C, E; Fig. S3C**). In contrast, DP and K8 displayed distinct recruitment kinetics, whereby both proteins exhibited continued junctional accumulation between 5 and 15 min, with DP increasing from 2.1-fold at 5 minutes to 4.1-fold at 15 min (a 2-fold increase), and K8 increased from 1.3-fold to 2.2-fold (a 1.6-fold increase). This was followed by a subsequent decline in junctional levels, which was evident at 30 min (DP: 1.5-fold; K8: 1.7-fold) (**Fig. 2D, F; Fig. S3D**). Overall, these results reveal two distinct recruitment behaviours among junctional complexes with respect to stress duration: stable responders with sustained levels (ZO-1, E-cad, PG) and adaptive responders (DP and K8), showing time-dependent accumulation (**Fig. 2E, F**).

**Figure 2.**
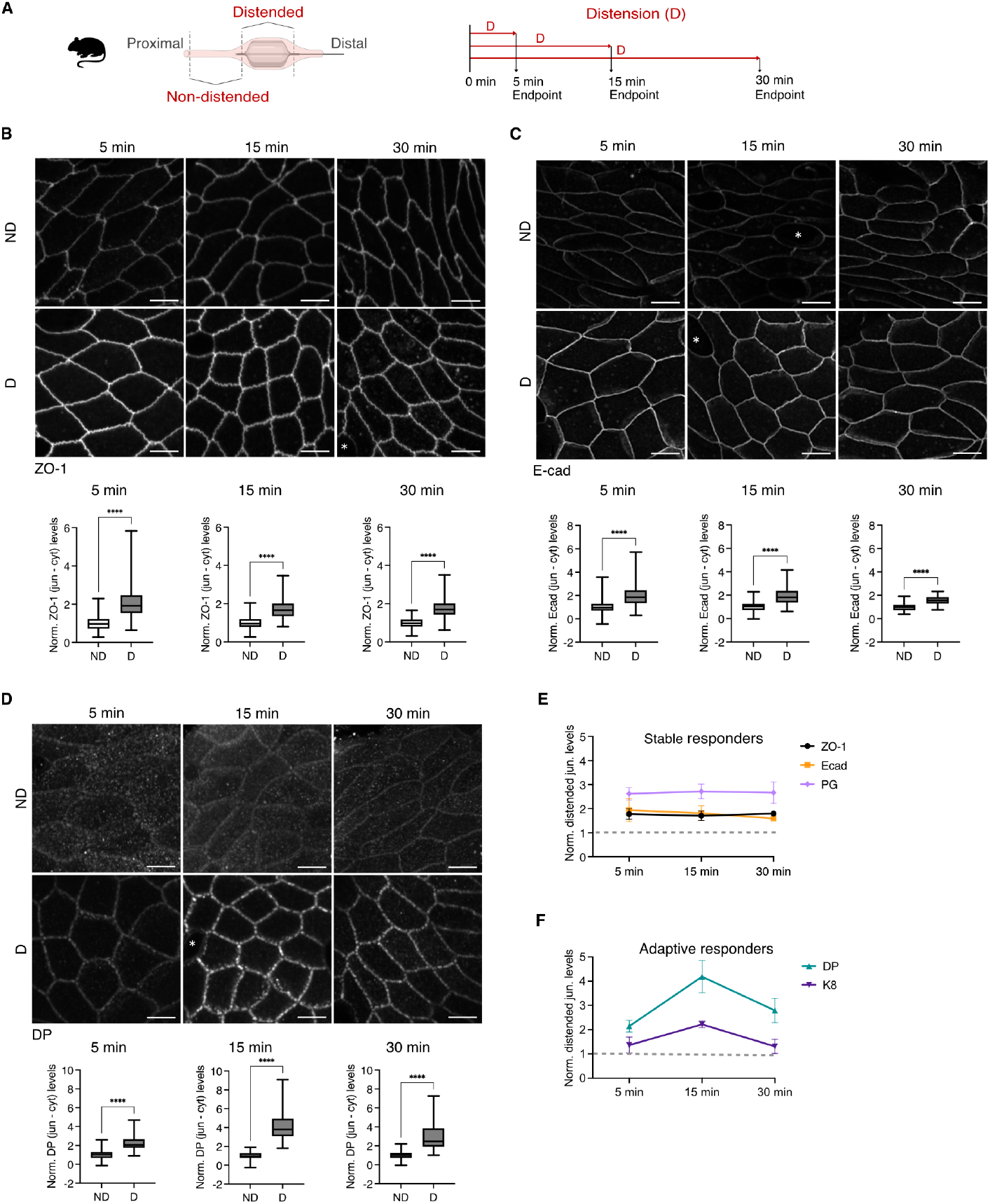
Time-resolved distension reveals distinct junctional recruitment dynamics. **(A)** Scheme showing experimental approach for catheter-mediated *in vivo* distension of the mouse colon. **(B)** Top: Representative *en face* images of colonic epithelium, stained for ZO-1 in non-distended (ND) and catheter-mediated distended (D) regions after 5,15 and 30 min of distension. Maximum Z-projections (1-3 µm range). Scale bar: 5 µm. Bottom (left to right): Box plots showing junctional ZO-1 intensity in ND and D regions after 5, 15 and 30 min of distension (n=26-125 junctions/condition, N=4-7 independent experiments). **(C)** Top: Representative *en face* images of colonic epithelium, stained for E-cad in ND and D regions after 5,15 and 30 min of distension. Maximum Z-projections (1-3 µm range). Scale bar: 5 µm. Bottom (left to right): Box plots showing junctional E-cad intensity in ND and D regions after 5, 15 and 30 min of distension (n=23-64 junctions/condition, N=2-4 independent experiments). **(D)** Top: Representative *en face* images of colonic epithelium, stained for DP in ND and D regions after 5,15 and 30 min of distension. Maximum Z-projections (1-3 µm range). Scale bar: 5 µm. Bottom (left to right): Box plots showing junctional intensity for DP in ND and D regions after 5, 15 and 30 min of distension (n=21-65 junctions/condition, N=3-6 independent experiments). **(E)** Comparative line plot showing junctional intensity fold change for ZO-1, E-cad and PG after 5, 15 and 30 min of distension. Dotted line represents the normalised average for ND. **(F)** Comparative line plot showing junctional intensity fold change for DP and KRT8 after 5, 15 and 30 min of distension. Dotted line represents the normalised average for ND. Statistical analysis was performed using the Mann-Whitney U test. Significance is denoted as: ****p < 0.0001, ***p < 0.0005, ** p < 0.005, ns: non-significant. ND: Non-distended, D: Catheter-mediated distended. *: goblet cell.

### 3. Perijunctional myosin II accumulates during distension and is required for junctional remodelling *in vivo*

Upon controlled distension, we observed a transient reduction in apical cell area (22%) (**Fig. S4A, B**). Since apical constriction is typically driven by actomyosin contractility, this suggests activation of myosin II in response to colonic distension. We thus decided to investigate the potential role of non-muscle myosin II (NMMII), a key cytoskeletal motor involved in epithelial remodelling under force. NMMIIA, the predominant non-muscle myosin II isoform localized at the perijunctional actomyosin belts, is well characterised for its role in generating contractile forces (*24, 25*). To examine the kinetics of perijunctional NMMIIA in distended versus non-distended tissues, we performed catheter-mediated *in vivo* distension, as before, in mice expressing GFP-tagged endogenous NMMIIA (*24*). NMMIIA showed pronounced enrichment at junctions during distension, peaking at 15 min (2.6-fold increase compared to 5 min) and coinciding with the transient reduction in apical cell area (**Fig. 3A, B**), suggesting activation of myosin II–mediated contractility. Notably, we also confirmed the recruitment of NMMIIA at the perijunctional belt in the faeces-distended part of the colon (**Fig. S4C, D**).

**Fig. 3.**
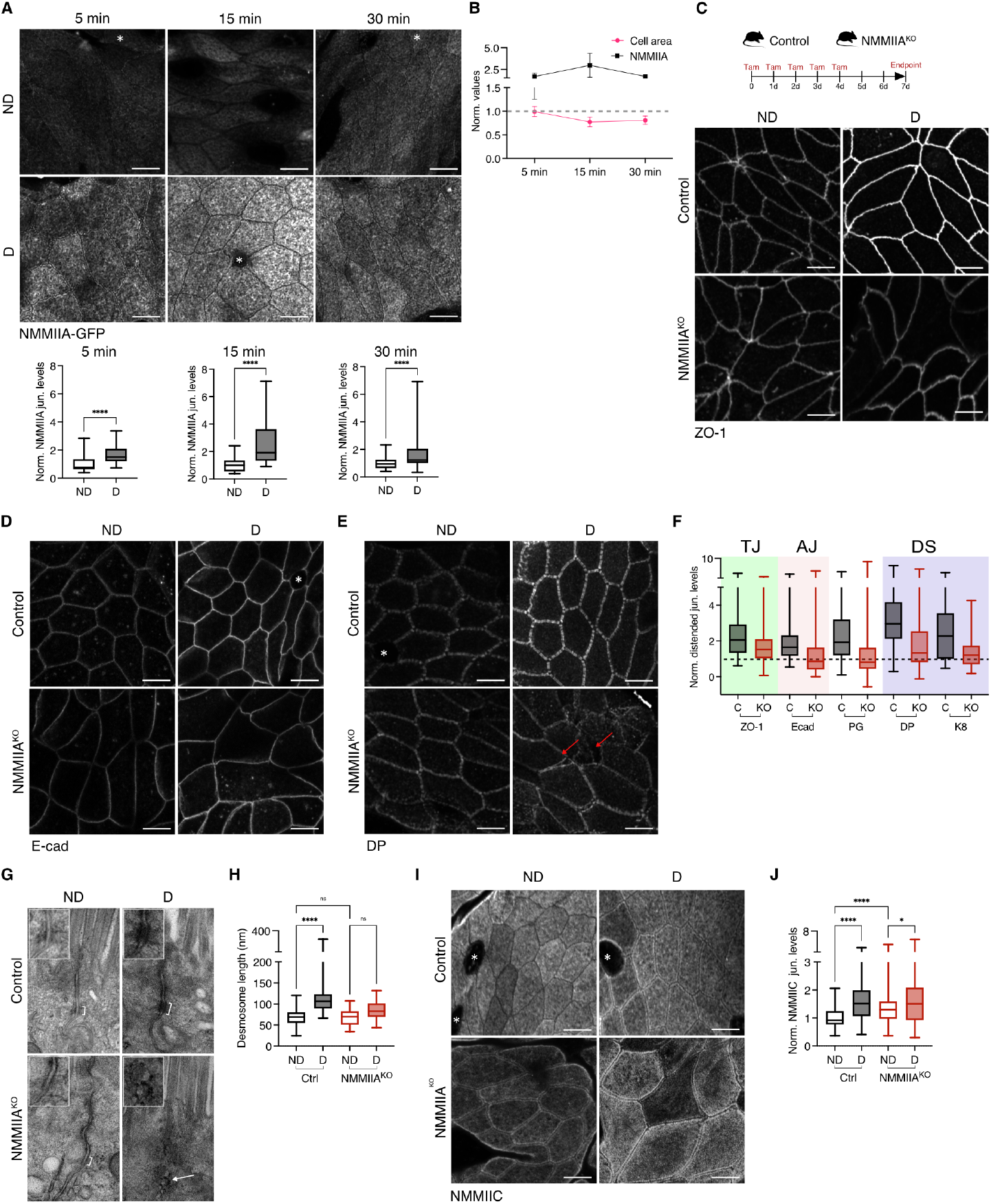
Junctional remodelling during colonic distension requires non-muscle myosin IIA. **(A)** Top: Representative *en face* images of colonic epithelium showing NMMIIA-GFP in ND and D regions after 5, 15 and 30 min of distension. Maximum Z-projection (1-3 µm range). Scale bar: 5 µm; Bottom (Left to right): Box plots showing junctional NMMIIA-GFP intensity in distended in ND and D regions after 5, 15 and 30 min of distension (n=35-62 junctions/condition, N=2 independent experiments. **(B)** Line plot showing mean fold change in cell area and NMMIIA-GFP intensity after 5, 15 and 30 min of distension. Dotted line represents the normalised average for ND. **(C)** Top: Scheme of the experimental procedure: Villin-CreERT2; Myh9^fl/fl^ mice were injected intraperitoneally (i.p.) for 5 consecutive days with tamoxifen and sacrificed 3 days after the last injection (endpoint). Villin-CreERT2^-/-^ and Villin-CreERT2^+/-^ animals were used as control and knockout (NMMIIA^KO^), respectively; Tam, tamoxifen. Bottom: Representative image of colonic epithelium stained for ZO-1 in control and NMMIIA^KO^ mice from ND and faeces-D regions. Maximum Z-projection (1-3 µm range). Scale bar: 5 µm. **(D-E)** Representative images of colonic epithelium stained for E-cad (D) and DP (E) in control and NMMIIA^KO^ mice from ND and faeces-D regions. Maximum Z-projection (1-3 µm range). Red arrow represents fractured DP junctions. Scale bar: 5 µm. **(F)** Comparative box plot showing fold change in junctional protein levels (tight junctions (TJ), adherens junctions (AJ) and desmosomes (DS)) in faeces-D regions of controls and NMMIIA^KO^ mice. Dotted line represents the normalised average for ND **(G)** Representative TEM images showing desmosomes (white bracket or arrow; enlarged insets at top left corner) in control and NMMIIA^KO^ mice from ND and faeces-D region. Scale bar: 100 nm. **(H)** Box plot showing desmosome length in control and NMMIIA^KO^ mice from ND and faeces-D regions (n=10-27 junctions/condition, N=2 independent experiments). **(I)** Representative image of colonic epithelium stained for NMMIIC in control and NMMIIA^KO^ mice from ND and faeces-D regions. Maximum Z-projection (1-3 µm range). Scale bar: 5 µm. **(J)** Box plots showing junctional intensity levels in ND and D for NMMIIC in control and NMMIIA^KO^ (n=30-60 junctions/condition, N=4 independent experiments). Statistical analyses were performed using the Mann–Whitney U test (A) or Kruskal-Wallis test followed by Dunn’s multiple comparison (H, J). Significance is denoted as: ****p < 0.0001, ***p < 0.0005, ** p < 0.005, ns: non-significant. ND: Non-distended, D: Distended. *: goblet cell.

To investigate the role of NMMIIA in epithelial junctional remodelling in response to distension, we used an inducible and gut epithelium-specific NMMIIA knockout (NMMIIA-KO) mouse model (**Fig. 3E**), which is indistinguishable from controls in terms of body weight and faecal size and morphology (**Fig. S4E**). Upon NMMIIA depletion, we observed a significant reduction in the faecal distension-dependent recruitment of multiple markers at all three types of cell-cell junctions, including the desmosomes with associated keratin 8 filaments (**Fig. 3E-H; Fig. S4F-H, K-N**). TEM imaging further revealed disrupted desmosomal plaques organization in distended NMMIIA-KO tissues (**Fig. 3G, H**). Notably, the apical cell area, already elevated in non-distended NMMIIA-KO tissue, increased further upon distension, exceeding the levels observed in controls (**Fig. S4I**), indicating that loss of NMIIA removes contractile restraint, rendering cells more susceptible to strain. Importantly, we observed no reduction in the total levels of junctional proteins in NMMIIA-KO mice compared to controls, indicating that NMMIIA promotes junctional complex accumulation through recruitment rather than by regulating their expression (**Fig. S5A, B**). Despite being significantly reduced, junctional recruitment in response to distension was not fully abolished in NMMIIA-KO. We thus investigated whether other myosin II isoforms could compensate. Of the other two isoforms – NMMIIB and NMMIIC – NMMIIB is not expressed in the gut epithelium (*25*), while NMMIIC localizes to the terminal web, a cortical actin-rich zone distinct from the perijunctional actomyosin belts (*26*). Interestingly, upon distension in wild-type mice, NMMIIC was also recruited to the perijunctional belt (**Fig. S5C, D**). Unlike NMMIIA, for which levels peaked at 15 min, NMMIIC levels remained constant across all time points. Interestingly, in non-distended conditions in NMMIIA-KO mice, NMMIIC was already localised to the junctions; upon distension, we observed an additional, significant increase (**Fig. 3I, J**). Total NMMIIC protein levels were unchanged in NMMIIA-KO mice (**Fig. S5A, B**), suggesting that NMMIIC is translocating from the terminal web to the junctions and could thus potentially compensate for NMMIIA depletion. Altogether, our data suggest that NMMIIA is actively recruited to junctions upon distension and that its presence is important for mediating the accumulation of junctional complexes in response to mechanical stress *in vivo*.

### 4. Junctional recruitment in response to mechanical distension depends on myosin II activation mediated by mechanosensitive ion channels

To further explore the mechanism of force-mediated junctional recruitment, we turned to pharmacological manipulation (**Fig. 4A**) and adapted our *ex vivo* gut explant culture system (*27*) to colon tissue. Explants from non-distended regions of the colon were maintained at an air-liquid interface and subjected to defined compression from above using a fixed weight, to mimic colonic distension (**Fig. 4B; Fig. S6A, B**). Recapitulating the *in vivo* system, distension of control explants induced robust accumulation of junctional complexes and, importantly, triggered phosphorylation of the myosin regulatory light chain (MLC), marking activation of contractility under mechanical stress (**Fig. 4C-F; Fig. S6; Fig. S7**). To target all myosin II isoforms simultaneously, we used blebbistatin, a general myosin II inhibitor known to reduce cortical contractility (*28*). Myosin II inhibition completely abolished the recruitment response across all three types of junctions, resulting in aberrant cell and junctional morphology (**Fig. 4C, F; Fig. S6C-G**). To investigate the mechanism underlying the activation of myosin II upon distension, we examined two key pathways: Rho-GTPase-mediated activation of ROCK, and calcium-calmodulin-dependent activation of MLCK. Both pathways converge on myosin II by phosphorylating the regulatory light chain (MLC), thereby enhancing contractility. Pharmacological inhibition of either ROCK (Y27632) or MLCK (ML7) blocked MLC phosphorylation and fully suppressed junctional complex recruitment under distension (**Fig. 4F; S6H-O; Fig. S7A-H**). Conversely, increasing myosin II activity by inhibiting MLC phosphatase with Calyculin A led to enhanced phosphorylation of MLC and increased junctional recruitment (ZO-1, 1.5-fold; pMLC, 1.7-fold), compared to mechanical distension alone (**Fig. 4D-F; Fig. S7I**). Collectively, these data suggest that myosin II activation via both ROCK and MLCK pathways is critical for the recruitment of junctional complexes upon distension. Moreover, the fact that weight-mediated distension of colonic explants *ex vivo*, which lack luminal contents and muscle contractility, still induced junctional recruitment, remarkably indicates that this response is epithelium-autonomous and directly triggered by mechanical forces rather than by chemical cues from faeces (*29, 30*) or muscle contractility.

**Fig. 4.**
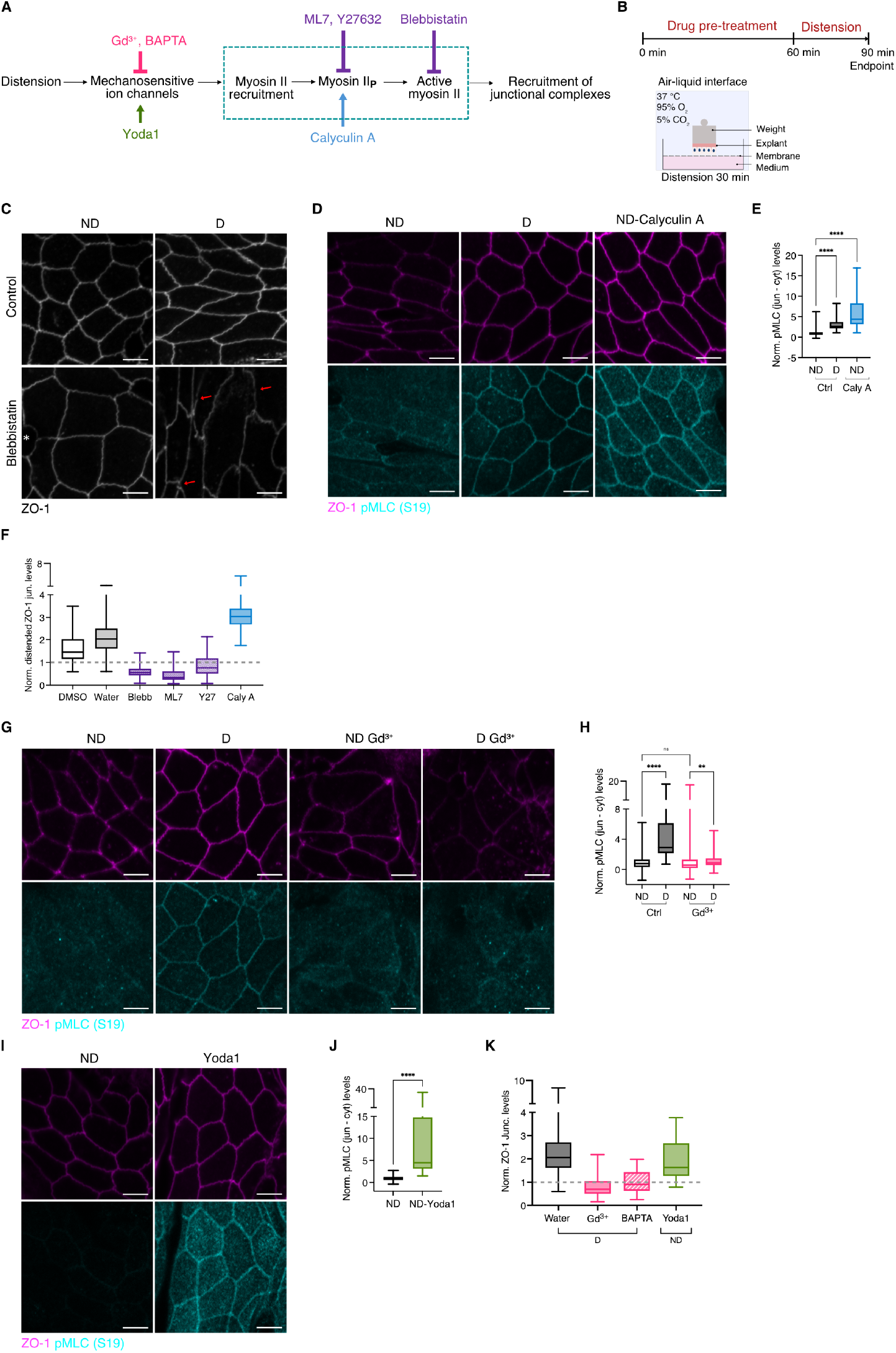
Myosin II activity regulated by mechanosensitive calcium influx is essential for junctional recruitment upon distension. **(A)** Scheme showing the potential mechano-adaptation pathway in response to distension in the colonic epithelium, indicating where each inhibitor or activator acts within the pathway. **(B)** Scheme depicting experimental approach for *ex vivo* culture and distension of colonic explants. **(C)** Representative image showing colonic explants, stained for ZO-1 in ND and D after blebbistatin or vehicle control (DMSO) treatment, as indicated in (B). Maximum Z-projection (1-3 µm range). Red arrows represent discontinuous ZO-1 staining. Scale bar: 5 µm. **(D)** Representative image showing colonic explants stained for ZO-1 (magenta) and pMLC (S19) (cyan) after Calyculin A or vehicle control (water) treatment (ND and D), as indicated in (B). Maximum Z-projection (1-3 µm range). Scale bar: 5 µm. **(E)** Box plot showing junctional pMLC intensity in Calyculin A- and vehicle control (water)-treated ND and D explants (n=53-59 junctions/condition, N=3 independent experiments). **(F)** Comparative box plot showing junctional ZO-1 intensity fold change in distended explants after pre-treatment with upstream myosin II inhibitors or Calyculin A. Dotted line represents the normalised average for ND. **(G)** Representative image showing colonic explants, stained for ZO-1 in ND and D after Gd^3+^ or vehicle control (water) treatment as indicated in (B). Maximum Z-projection (1-3 µm range). Scale bar: 5 µm). **(H)** Box plot showing junctional pMLC intensity in Gd^3+^ treated and vehicle control (water)-treated ND and D explants (n=58-61 junctions/condition, N=5 independent experiments). **(I)** Representative images showing colonic explants, stained for ZO-1 (magenta) and pMLC (S19) (cyan) in ND after Yoda1 or vehicle control (DMSO) treatment for 15 min. Maximum Z-projection (1-3 µm range). Scale bar: 5 µm. **(J)** Box plot showing junctional pMLC intensity in Yoda1-treated and vehicle control (DMSO)-treated ND explants (n=53-59 junctions/condition, N=2 independent experiments **(K)** Comparative box plot showing junctional ZO-1 intensity fold change in explants after pre-treatment with mechanosensitive ion channel inhibitors or Piezo1 activator. Dotted line represents the normalised average for ND. Statistical analyses were performed using the Mann-Whitney U test (J) or Kruskal-Wallis test followed by Dunn’s multiple comparison (F) and (H). Significance is denoted as: ****p < 0.0001, ***p < 0.0005, ** p < 0.005, ns: non-significant. ND: Non-distended, D: Distended. *: goblet cell.

To investigate whether the observed myosin II recruitment and MLC phosphorylation are mediated via calcium influx upon distension, we inhibited the activity of mechanosensitive ion channels either via gadolinium hexahydrate chloride (Gd^3+^) (*31*) or blocked calcium entry by chelating extracellular calcium with BAPTA (*32*). These treatments abolished both the MLC phosphorylation and the junctional recruitment in response to distension (**Fig. 4G-H, K; Fig. S7J-N**). In agreement, activation of the mechanosensitive calcium channel Piezo-1 using the chemical activator Yoda1 (*33*) resulted in myosin II activation and recruitment, as well as the accumulation of junctional ZO-1 (2.2 fold; **Fig. 4I-K; Fig. S7O-P**). Taken together, our findings establish myosin II activation as essential for junctional recruitment in response to mechanical distension, and identify calcium influx through the Piezo1 ion channel as a potent and sufficient upstream trigger of this contractile response.

### 5. Junctional reinforcement protects the colonic barrier from mechanical stress-induced breach

During homeostasis, epithelial barrier integrity is maintained by adhesive junctional complexes, primarily tight junctions (TJs), which form a continuous belt-like structure around cells, sealing the paracellular space and preventing luminal translocation. Since we showed that junctional recruitment under mechanical stress is impaired when myosin II is inactivated, we next asked whether barrier integrity is also compromised under these conditions. In addition to impaired junctional recruitment, distended regions in NMMIIA-KO tissue and blebbistatin-treated explants exhibited irregular and discontinuous ZO-1 staining (**Fig. 3C; S4J; Fig. 4C**), suggesting a loss of TJ integrity. To determine whether such discontinuous junctions represent a breach in the epithelial barrier (**Fig. 5A**), we assessed luminal accessibility of E-cad using a sequential staining assay. Antibodies against the extracellular domain of E-cadherin were applied to intact, freshly fixed tissue prior to permeabilization; luminal antibody binding and subsequent E-cad labelling indicate that tight junctions were breached (*34*) (**Fig. 5B**). While we rarely detected “accessible E-cad”-positive junctions upon distension in control mice, NMMIIA-KO showed a significant increase in breached junctions under distension (Control: 5%; NMMIIA-KO: 57%; **Fig. 5C, D**). A similar trend of increased barrier breach in distended colonic explants was also observed upon myosin II inhibition (**Fig. 5E, F, H; Fig S8A, B**). Additionally, inhibition of mechanosensitive ion channels using gadolinium salts induced barrier defects upon distension (**Fig. 5G, H**). These findings indicate that loss of NMMII function – through abolished MLC phosphorylation and/or inhibited calcium influx through mechanosensitive ion channels – compromises barrier integrity under extrinsic mechanical stress. Such compromised junctions may allow translocation of luminal contents, including microbial metabolites and toxins, potentially triggering inflammation and gut disorders. Notably, in some distended regions of NMMIIA-KO tissue, we observed junctional fractures accompanied by bacteria in the paracellular space (**Fig. S8C**).

**Fig. 5.**
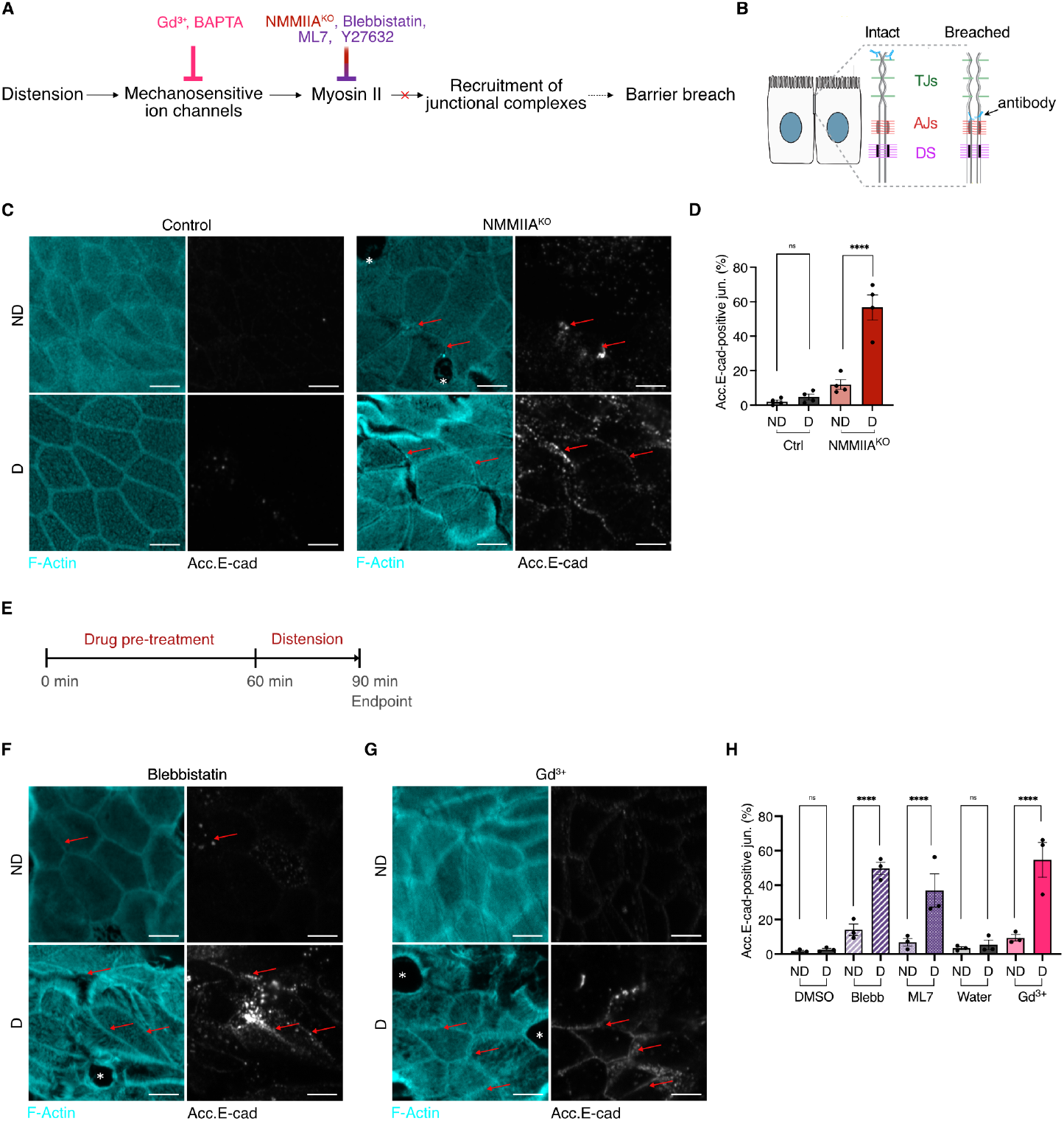
Junctional recruitment protects epithelial barrier integrity during distension. **(A)** Schematic of the proposed mechanoadaptation pathway in colonic epithelium, illustrating how failed junctional recruitment may lead to barrier breach. **(B)** Scheme illustrating the accessible E-cad assay (see Materials and Methods), where E-cad antibody stains only if the tight junctions are breached. **(C)** Representative images of colonic epithelium stained for F-actin (cyan) and acc. E-cad (grey) in control and NMMIIA^KO^ mice from ND and faeces-D regions. Red arrows indicate breached junctions. Maximum Z-projection (1-3 µm range). Scale bar: 5 µm. **(D)** Bar plot showing percentage of accessible E-cad (acc. E-cad)-positive junctions in control and NMMIIA^KO^ mice from ND and faeces-D regions (n=19-57 junctions/condition, N=4 independent experiments). **(E)** Scheme showing experimental approach for accessible E-cad assay in *ex vivo* culture. **(F)** Representative images showing colonic explants stained for acc. E-cad in ND and D after treatment with blebbistatin as indicated in (E). Maximum apical Z-projection (1-3 µm range). Scale bar: 5 µm. **(G)** Representative images showing colonic explants stained for acc. E-cad in ND and D after treatment with Gd^3+^ as indicated in (E). Maximum Z-projection (1-3 µm range). Scale bar: 5 µm. **(H)** Comparative bar plot showing percentage of positive junctions for acc. E-cad in ND and D explants after treatment with vehicles (DMSO and water), Blebbistatin, ML7 and Gd^3+^ as indicated in (E) (n>200 junctions/condition, N=3 independent experiments). Statistical analysis was performed using the one-way ANOVA followed by Bonferroni post hoc test (D) and (H). Bar plots: whiskers denote SEM. Significance is denoted as: ****p < 0.0001, ***p < 0.0005, ** p < 0.005, ns: non-significant. ND: Non-distended, D: Distended. Red arrows indicate breached junctions. *: goblet cell

In conclusion, we show that colonic mucosa undergoes coordinated cell- and tissue-scale remodelling to accommodate faeces, with junctional complexes responding by recruiting their constituent proteins to reinforce the barrier. Our findings highlight NMMII as a key mediator of this recruitment, activated by calcium influx through mechanosensitive ion channels. Loss of junctional recruitment response leads to barrier defects under mechanical stress, underscoring the critical role of junctional reinforcement in maintaining gut epithelial integrity.

## Discussion

Few studies have investigated how different adhesive junctions interact under mechanical stress, and these have mostly relied on loss-of-function experiments in *in vitro* models (*29, 35*) or early embryos (*45*). Here, we report that during physiological faeces-mediated colon distension in adult mice, epithelial cells experience mechanical forces leading to recruitment of proteins from all three types of junctional complexes (TJs, AJs, and desmosomes). Importantly, we have for the first time analysed the coordination and temporal regulation of junctional recruitment at all junction types. This revealed two distinct classes of junctional responders to mechanical force. The stable responders – including ZO-1, E-cad, and plakoglobin – were recruited within 5 min of distension and remained at (increased) constant levels for up to 30 min, suggesting a stable mechanoadaptive response. Given that ZO-1 is a core scaffolding protein of tight junctions, its early and stable recruitment is likely critical in preserving barrier integrity under mechanical stress. Interestingly, plakoglobin – which can bind to both AJs and desmosomes – has previously been identified as a mechanosensitive protein in Xenopus mesendoderm cells, where it plays a key role in the reorganization of keratin intermediate filaments toward sites of mechanical stress (*36*). Therefore, the sustained levels of plakoglobin during distension in the gut epithelium may support both AJ and desmosome assembly, enhancing tissue mechanical resilience together with ZO-1 at TJ. In contrast, adaptive responders (desmoplakin and keratin 8) showed progressive accumulation, peaking around 15 min of distension, with higher fold changes than those observed for TJ and AJ proteins. This observation is consistent with a recent in vitro study demonstrating that intermediate filaments undergo strain-stiffening under prolonged mechanical load, characterized by increased bundling of keratin 8, to provide structural reinforcement against sustained mechanical force (*37*). Beyond our comparative analysis of all three junctional types *in vivo*, our results on plakoglobin, which can associate with both AJs and desmosomes (*36, 38*), may contribute to both sustained and adaptive junctional responses, suggesting a key role as a mediator of junctional crosstalk in response to mechanical force (*39, 40*). We further showed that NMMII activity governs mechanoadaptation of all adhesive junctions in the gut epithelium, including desmosomes and their associated intermediate filaments, suggesting that NMMII acts early as a central regulator – or “hub” – of epithelial adhesion, potentially facilitating crosstalk between different junctional complexes. We observed accumulation and activation of NMMII in the perijunctional space during colonic distension; similar phenomena have been described at junctions during Drosophila development, where stimulation of contractility promotes myosin recruitment and actomyosin remodelling (*41, 42*). The gut epithelium expresses two relevant NMMII isoforms: NMMIIA, predominantly associated with AJs and exhibiting a low duty ratio that enables dynamic contractile force generation, and NMMIIC, which localises to the terminal web and regulates microvillar architecture (*25*). Upon distension, NMMIIA gradually accumulated at junctions, peaking at 15 min, coinciding with maximal transient apical constriction and the recruitment of adaptive responders, suggesting that NMMIIA-mediated contractility drives their progressive junctional accumulation. Interestingly, NMMIIC also relocalised to junctions in response to force; however, unlike NMMIIA, its levels remained relatively constant throughout the distension period. Loss of NMMIIA impaired junctional recruitment, indicating that its presence is a prerequisite for recruitment of both stable and adaptive responders. Notably, in NMMIIA-KO tissue, NMMIIC relocalised to junctions even under non-distended conditions, suggesting a potential compensatory role. To simultaneously target both isoforms, we treated colon explants with pharmacological inhibitors of myosin II activity. The complete loss of junctional recruitment following broad myosin II inhibition in these *ex vivo* experiments suggests that NMMIIC has a supportive role in this process. Inhibition of upstream myosin pathways – either via the Rho-ROCK pathway or MLCK – was sufficient to abolish recruitment across all three types of adhesive junctions upon force, while increasing myosin activity (using calyculin A) enhanced recruitment of junctional proteins, even in the absence of applied force. Our results suggest that these pathways synergise in an essential and non-redundant manner to trigger NMMIIA-mediated mechanotransduction. We showed that this effect is potentially mediated by calcium signalling, which is known to promote myosin contractility through activation of both RhoA signalling and calmodulin/MLCK pathway (*7, 43*–*45*). Calcium signalling in epithelial cells is a key mediator of mechanotransduction, converting mechanical forces into biochemical signals via mechanosensitive ion channels. A recent study of global Piezo knockout in the gut epithelium *in vivo* reported severe defects in stem cell differentiation, resulting in diarrhoea, weight loss, and early lethality (*46*). However, the more immediate role of Piezo-mediated calcium influx in junctional dynamics and barrier integrity had not been addressed. Here, we used acute pharmacological modulation of mechanosensitive calcium influx *ex vivo*, including selective activation of Piezo1, enabling us to probe short-term effects on epithelial junctions and barrier function. Using colonic explants, inhibition of mechanosensitive ion channels with gadolinium salts resulted in loss of MLC phosphorylation and abolished ZO-1 recruitment. As gadolinium broadly targets multiple mechanosensitive ion channels, including Piezo1 and TRPV (TRPV1 and TRPV4) channels (*47*), we further investigated the specific role of Piezo1 using its chemical activator Yoda1. Chemical activation of Piezo1 in non-distended explants significantly increased MLC phosphorylation and ZO-1 localization at junctions, indicating that Piezo1 activation is sufficient to trigger myosin II recruitment and activation, an early mechanosensitive event, and to drive recruitment of junctional complexes.

Remarkably, the “reinforced” junctional state was strictly dependent on continuous extrinsic mechanical load: upon its removal, the junctional levels rapidly reverted to baseline levels, indicating that sustained force is necessary for the recruitment and stabilisation of junctional complexes. Such dynamic regulation may arise from tension-induced conformational changes in junctional components (*9, 48, 49*) or from force-mediated, phosphorylation-dependent mechanisms, as the phosphorylation states of plakoglobin and desmoplakin are known to regulate desmosome assembly and disassembly (*50, 51, 12*). The rapid reversion of junctional levels to baseline after force removal in the gut epithelium suggests a mechanism to conserve energy, since maintaining junctions in a reinforced state during distension is energetically demanding, relying on ATP-dependent processes such as actin polymerization, actomyosin contractility, protein trafficking, and signalling (*52*–*54*).

Finally, our findings establish junctional recruitment as a mechano-adaptive response to mechanical stress, essential for preserving barrier integrity. This mechanism may be particularly relevant in gut pathologies such as toxic megacolon, severe constipation, luminal obstruction by tumours, and fibrotic strictures in inflammatory bowel disease (IBD), where excessive mechanical distension could worsen disease outcomes (*55*–*57*). Consistently, studies in a gut epithelium-specific constitutive NMMIIA knockout mouse model showed increased intestinal permeability under basal conditions, further exacerbated by experimental colitis, highlighting a protective role for NMMII in maintaining epithelial integrity (*58*). In conclusion, we demonstrate that the adult colonic epithelium mounts a robust mechano-adaptive response to mechanical stress, whereby NMMII, activated by mechanosensitive calcium influx, coordinates junctional reinforcement to preserve barrier integrity. This reveals a fundamental mechanism by which the gut epithelium sustains homeostasis under physiological mechanical stress.

## Acknowledgments

We thank B. Bénazéraf, E. Theveneau, C. Bierkamp, A. Merdes, K. Ancelin, D. Pinheiro, R. Galupa and all members of the Krndija lab for their critical feedback and suggestions. We acknowledge the support and contribution of the ANEXPLO mouse facility, the Multiscale Electron Imaging (METI) facility and the Light Imaging Toulouse CBI (LITC) facility at the CBI. This work was supported by an ATIP-Avenir grant to D.K. from CNRS, and by fellowships to V.K. from the École Doctorale Biologie, Santé, Biotechnologies (BSB) and the Fondation pour la Recherche Médicale (FRM).

## Author contributions

Conceptualization: D.K.; Methodology: V.K., N.C., D.K.; Investigation: V.K., A.F., S.B.F., J.C.; Project administration: V.K., D.K.; Supervision: D.K.; Funding acquisition: D.K.; Writing – original draft: V.K., D.K.; Writing – review & editing: V.K., J.C., D.K.

## Competing interests

The authors declare no competing interests.

## Materials & Correspondence

Correspondence and requests for materials should be addressed to D.K

## Notes

### Competing Interest Statement

The authors have declared no competing interest.

### Summary of Updates

Mistakes in the text and missing references

