## Supplementary Material for "Robust pan-junctional reinforcement preserves the gut epithelial barrier under mechanical stress"

**The PDF file includes:**

Materials and Methods
Figs. S1 to S8
References

### Materials and methods

#### Mice

All mouse husbandry and handling procedures were approved and conducted in accordance with European and National Regulations (Protocol authorization APAFIS #38994-2023062718345638, APAFIS #14513-2018040313435341 v6.). Mice were housed in the Specific and Opportunistic Pathogen-Free (SOPF) facility at the CBI and CREFRE SOPF facility of INSERM Purpan. Experiments were performed on age-matched littermates (8–12 weeks old) of both sexes, as no phenotypic differences were observed between males and females. C57BL/6N mice were obtained from Charles River Laboratories (France). Villin: $Cre^{ERT2}$ mice were crossed with $Myh9^{flox}$ (Mutant Mouse Resource and Research Center, stock 032096-UNC) to generate $NMIIA^{KO}$ mice. Villin: $Cre^{ERT2}$ and $NMIIA^{GFP}$ mice were kindly provided by Dr. D. Matić Vignjević (Institut Curie).

#### Tamoxifen treatment

To induce intestinal epithelium-specific knockout of Myh9 (NMIIA), Villin: $Cre^{ERT2}/Myh9^{fl/fl}$ and $Myh9^{fl/fl\wedge}$ control mice were injected with 100 $\mu$L of tamoxifen (50 mg/kg; Euromedex SE-S1238) intraperitoneally (i.p.) once daily for five consecutive days and culled three days after the last injection.

#### Antibodies

For immunofluorescence, we used the following antibodies: rat anti-ZO-1 (Millipore, MABT11) at 1:200, rat anti-E-cadherin (Invitrogen, 1319800) at 1:200, rabbit anti-occludin (Proteintech, 27260-1-AP) at 1:50, mouse anti-α-catenin (Santa Cruz, SC-9988) at 1:50, mouse anti-β-catenin (Santa Cruz, SC-7963) at 1:50, rabbit anti-NMMIIC (Proteintech, 909802) at 1:100, rabbit anti-pMLC (Ser19) (Cell Signalling, 3671) at 1:100, sheep anti-desmocollin-2 ( Biotechne, AF7490) at 1:50), mouse anti-γ-catenin (Plakoglobin) (Santa Cruz, SC-514115) at 1:50, mouse anti-desmoplakin1/2 (Progen, 651109) at 1:50, rabbit anti-cytokeratin 8 (Abcam, ab240986) at 1:200, rabbit anti-laminin (Merck, L9393) at 1:200. All the secondary antibodies (Molecular Probes) were used at a 1:200 dilution. F-actin was labelled with Rhodamine- or Alexa Fluor 647-conjugated phalloidin (Invitrogen, R415 and A30107) at 1:200. DNA was stained using DAPI (Sigma, D9452) at 5 $\mu$g/mL. For western blotting, the same antibodies as those listed for immunofluorescence were used. Additional antibodies specific to western blotting include: rabbit anti-ZO1 (Proteintech, 21773-1-AP ) at 1:10000, rabbit ,anti-NMMIIC (Cell signalling, 8189T) at 1:5000, rabbit anti-NMMIIA (Biolegend, 909802) at 1:5000, rabbit anti-cytokeratin 8 (Abcam, ab227644) at 1:1000, rabbit anti-GAPDH (Sigma, G9545) at 1:40000. HRP-conjugated secondary antibodies against rabbit, mouse and rat IgG (1:5000) were purchased from Jackson Immunoresearch.

**Tissue fixation and sectioning**

Mice were culled as indicated, and the entire colon were isolated and fixed immediately in 4% paraformaldehyde (Electron Microscopy Sciences) in PBS for 1 hour at room temperature (RT), without flushing the luminal contents. After fixation, tissues were rinsed three times with 1× PBS and then processed by either transversal sectioning to visualize the apicobasal axis of epithelial cells, or by hand cutting into small sheets to image the epithelium from the apical (luminal) side, from both distended and non-distended regions. For transverse sections, the fixed tissues were cut into 150-200 $\mu$m-thick slices using a vibratome (Leica; VT1200). To prepare small sheets, the colon was cut open longitudinally along its length and small rectangular sections (≈4 $mm^2$) were dissected and isolated.

**3D immunofluorescence**

The fixed tissue sections were permeabilized with 1% Triton X-100 (Sigma) in PBS for 1 hour at RT followed by incubation with primary antibodies in washing buffer (0.2% Triton X-100/PBS) overnight at RT under gentle agitation. After incubation, the samples were washed three times for 1 hour each with the washing buffer, then incubated with secondary antibody, DAPI and with or without phalloidin, in the washing buffer. Following three washes of one hour each, the sections were mounted on slides so that the epithelial surface/plateaus were positioned directly beneath the coverslip using mounting medium (Aqua-Poly/Mount, Polysciences), according to the manufacturer's instructions. This ensured comparable imaging depths across all samples, thereby minimising variability in fluorescence intensity measurements arising from differences in optical path length.

**Transmission electron microscopy**

The colon was isolated as described above and fixed without flushing the luminal contents in 2.5% glutaraldehyde and 2% paraformaldehyde in 0.1 M cacodylate buffer (pH 7.2) overnight at 4°C. Post-fixation was performed with 1% osmium tetroxide in cacodylate buffer for 1 hour at RT, followed by washes in cacodylate buffer. The tissue was embedded in 2% low-melting-point agarose and stained with 1% uranyl acetate. Samples were dehydrated through a graded ethanol series (25%, 50%, 70%, and 90% for 15 min each, followed by three 30 min washes in 100% ethanol). Infiltration was performed in increasing concentrations of Epon resin (EMS, Embed 812) diluted in ethanol (25%, 50%, 75% for 1 h each at room temperature), followed by two 2 h incubations in 100% Epon at 37°C. Samples were embedded in Epon resin and polymerized for 48 h at 60°C. Ultrathin sections (~150 nm) were prepared using a Leica Ultramicrotome UCT, mounted on 200-mesh Formvar-carbon-coated copper grids, and stained with Uranyless (EM-grade) and 3% Reynolds lead citrate (EM-grade). TEM imaging was performed on a JEOL JEM-1400 at 80 kV equipped with a Gatan Orius digital camera.

**Colon relaxation assay**

Colon was isolated as described earlier, and segmented into regions with faeces (distended) or without faeces (non-distended). Faecal pellets were either gently removed from distended fragments using forceps or left inside, after which all samples including non-distended, distended, and relaxed fragments (fragments after faecal removal) were incubated in Leibovitz's L-15 medium (Gibco, 11415064) at 37°C for 5 min, followed by fixation and processing as described in the 3D immunofluorescence staining protocol.

**Catheter-mediated *in vivo* distension**

Colonic distension was performed as previously described (*1*). Briefly, the mice were anaesthetized via i.p. injection of ketamine and xylazine (6.7 mg/Kg and 133 mg/Kg, respectively). A Fogarty Thru-Lumen Embolectomy Catheter (4F; Edwards Lifesciences) with a deflated balloon was inserted 3 cm proximally from the anus. The balloon was then inflated with 1 ml water to a pressure of 80 mmHg/10.6 kPa, which was maintained for either 5, 15 or 30 minutes. After each time point, mice were culled, and the colon containing the inflated balloon was carefully isolated and fixed in 4% paraformaldehyde for 1 h at RT. Following fixation, the tissue was processed for 3D immunofluorescence as described above.

**Explant culture and treatments**

Colonic explant cultures were performed with slight modification to previously described method (*2*). Briefly, mice were sacrificed, and non-distended part of the colon were isolated and opened up longitudinally. Small sheets of tissue (≈ 4 $mm^2$) were cut and placed on a hydrophilic PTFE membrane insert with pore size of 0.4 $\mu$m (Millipore, PICM0RG50), epithelium facing the membrane. The explant medium consists of DMEM-F12 (Gibco, 12634010) supplemented with 2.5% FBS, 1X Glutamax (Fisher Scientific, 11574466), 1X Insulin-Transferrin-Selenium solution (Gibco, 41400045), 100 $\mu$g/mL transferrin (Santa Cruz, 391098), 10 ng/mL murine EGF (Peprotech, 315-09) and 1% anti-anti (Gibco, 15240062). Explants were cultured at 37°C in a humidified atmosphere of 95% $O_2$ and 5% $CO_2$, in the presence or absence of pharmacological inhibitors/chemical compounds.

To apply mechanical stimulation, tissues were either left non-distended or subjected to mechanical loading by placing a 2 g sterile, static weight directly on top of the explant's serosal side for 30 min (≈ 5 kPa for a 4 $mm^2$ surface area), adapting approaches previously used in epithelial mechanics (*3–5*). The estimated stress assumes uniform force distribution over the explant surface and does not correct for local deformation or tissue stiffness. Tissues were then fixed and processed for staining as described above.

For the combined pharmacological and mechanical treatments, the explants were pre-treated for 1 h with one of the following inhibitors or compounds: 170 $\mu$M blebbistatin (Selleckchem, S7099), 100 $\mu$M Y27632 (Y27), (Santa Cruz, sc-281642A), 100 $\mu$M ML-7 (Sigma, 475880), 100 $\mu$M calyculin-A (Abcam, ab141784), or 100 $\mu$M gadolinium (III) chloride ($Gd^{3+}$), (Sigma-Aldrich, G7532). Following pre-treatment, tissues were maintained under either non-distended or distended conditions for 30 min. For treatment with 500 μM Yoda1 (Sigma, SML1558), explants were incubated for 15 min in the non-distended condition. For 2 mM BAPTA (Millipore, 196418) treatment, the compound was added directly to the media, and samples were immediately subjected to distension or maintained non-distended for 30 min.

**Accessible E-cadherin assay**

The accessible E-cadherin assay was performed as previously described (*6*). Briefly, fixed tissue sections were incubated with a primary antibody against the extracellular domain of E-cadherin (ECCD-2) in PBS at RT under gentle agitation overnight. After three 1-hour washes in PBS, the samples were incubated with a secondary antibody in PBS overnight at RT, followed by a further three 1-hour washes. The tissues were then permeabilised with 1% Triton X-100 in PBS for one hour at RT and subsequently processed using the 3D immunofluorescence staining protocol described earlier.

**Western blotting**

The mouse colons were isolated and opened longitudinally. The distended regions containing faeces were either separated from the non-distended regions, or the entire colon was washed in PBS and cut into multiple fragments. The tissues were incubated in a solution of 30 mM EDTA and 1 mM DTT in PBS for 20 min on ice, followed by 30 mM EDTA/PBS for 15 min at 37°C. Samples were then vigorously agitated and centrifuged at 1200 rpm for 5 min at 4°C to isolate epithelial cells. Cells were lysed in RIPA buffer containing 50 mM Tris base, 150 mM NaCl, 1% SDS, 1% sodium deoxycholate, 1% Triton X-100, protease inhibitors (Roche, 11836170001) and phosphatase inhibitors I and II (Bimake, B15001, B15002). Samples were sonicated for 20 s and centrifuged at 12,000 × g for 10 min at 4°C. Protein concentration was measured using the DC assay (Bio-Rad, 5000112), and 25 μg of protein was mixed with 4× Laemmli buffer, boiled at 95°C for 5 min, and loaded onto 4–20% SDS–polyacrylamide gels (Bio-Rad, 4568095) under reducing conditions. Proteins were transferred to nitrocellulose membranes (Bio-Rad, 1704271), blocked with blocking buffer (Bio-Rad, 12010020) for 30 min, and incubated with primary antibodies for 1 h at room temperature or overnight at 4°C. After washing, membranes were incubated with HRP-conjugated secondary antibodies for 1 h at room temperature. Protein bands were revealed by ECL (chemiluminescence, Bio-Rad) and imaged with a ChemiDoc imaging system (Bio-Rad).

**Confocal 3D imaging**

All images were acquired from the plateau regions of the distal colon using transverse sections to visualize the apicobasal axis of epithelial cells, and sheet-like sections and explants to image the epithelium from the apical (luminal) side. Imaging was performed using a LSM 880 (Zeiss) inverted laser scanning microscope equipped with 405nm, 488nm, 561nm and 633nm lasers. Objectives used included 20x (z-step size: 0.8 $\mu$m), 40× (z-step size: 0.4 $\mu$m) oil immersion, and 63× (z-step size: 0.3 $\mu$m) oil immersion, with images captured at a resolution of 1024 × 1024 pixels.

**High resolution (Airyscan) 3D tissue imaging**

Images were acquired with LSM 880 (Zeiss) inverted laser scanning microscope equipped with Airyscan module. A 63× (z-step size: 0.2 $\mu$m) oil immersion objective was used and images were captured at a resolution of 1024x1024 resolution. Images were further processed using the Airyscan processing module in ZEN software.

**Image analysis**

All images were processed and analysed using ImageJ software (NIH). Optical Z-steps (0.3 µm intervals) were acquired to account for differences in cell height and tissue folding. Maximum intensity projections of image stacks (1–3 $\mu$m range) from *en face* sections, encompassing apical junctional complexes, were used for quantification. The Z-stack range for TJ and AJ proteins in each region of interest (ROI) was defined to encompass the perijunctional F-actin signal, ensuring that the F-actin belts of neighbouring cells were clearly distinguishable. Similarly, for desmosomes and associated proteins, the Z-stack range was defined to include the most-apical desmosome DP staining. Junctional fluorescence intensity was manually quantified using the line scan function by drawing a segmented line (10 pixels wide) orthogonal to randomly selected homotypic junctions, extending from vertex to vertex of the cell. To correct for cytoplasmic background, an equivalent line was drawn in the adjacent cytoplasm, and its intensity was subtracted from the junctional signal. For perijunctional proteins such as NMMIIA, NMMIIC, and keratin 8, intensity measurements were performed by line scans (10 pixels wide) encompassing the perijunctional regions between vertices. All intensity values were normalized to their respective non-distended controls within each experiment to calculate fold changes. Quantitative analysis of junctional and cytoplasmic protein distribution within cell was assessed using the Plot profile tool in image J. A line of constant length was drawn perpendicular to the junctions, extending into the cytoplasm, and average intensity was plotted along the normalized length.
Apical cell area was measured by segmentation on ZO-1 staining using Tissue Analyzer plugin in ImageJ and represented as area heatmap using Imaris software (Imaris 10.2; Oxford instruments).
Transverse images of colonic tissue were used to quantify cell and inter-cryptal aspect ratios. Cell aspect ratio was defined as the ratio of cell height to width in plateau regions, with F-actin staining used to demarcate cell boundaries and laminin to mark the basement membrane. Cell height was measured from the basement membrane to the tip of the microvilli, and width was measured as the distance between opposing lateral membranes at the cell midpoint. The inter-cryptal aspect ratio was calculated as the ratio of inter-cryptal region height to plateau width, with plateau width defined as the distance along the epithelial surface between adjacent crypt openings, and inter-cryptal height measured from the base of the inter-cryptal region to the epithelial surface.

**Statistics**

Statistical analysis was performed using GraphPad Prism 10.4.2. All experiments were independently repeated at least three times unless stated otherwise, and N represents the number of independent experiments. Normality of data was assessed using the Shapiro-Wilk test. For comparisons between two groups, normally distributed data were analysed using unpaired Student's t-tests, while non-normally distributed data were analysed using the Mann-Whitney test. For comparisons involving three or more groups, one-way ANOVA followed by Bonferroni's post hoc test was applied to normally distributed samples. For non-normally distributed samples, the Kruskal-Wallis test followed by Dunn's multiple comparison test was used. Data are presented as box-and-whisker plots – the box represents the interquartile range (25th–75th percentiles), the central line indicates the median, and the whiskers denote the minimum and maximum values. Bar charts represent average values from individual experiments and error bars indicate SEM. Statistical parameters, including sample size (n), statistical test performed, and significance values, are provided in the corresponding figure legends. Symbols used are: ns: $p > 0.05$; *: $p \leq 0.05$; **: $p \leq 0.01$; ***: $p \leq 0.001$; ****: $p \leq 0.0001$.

### Supplementary Figures

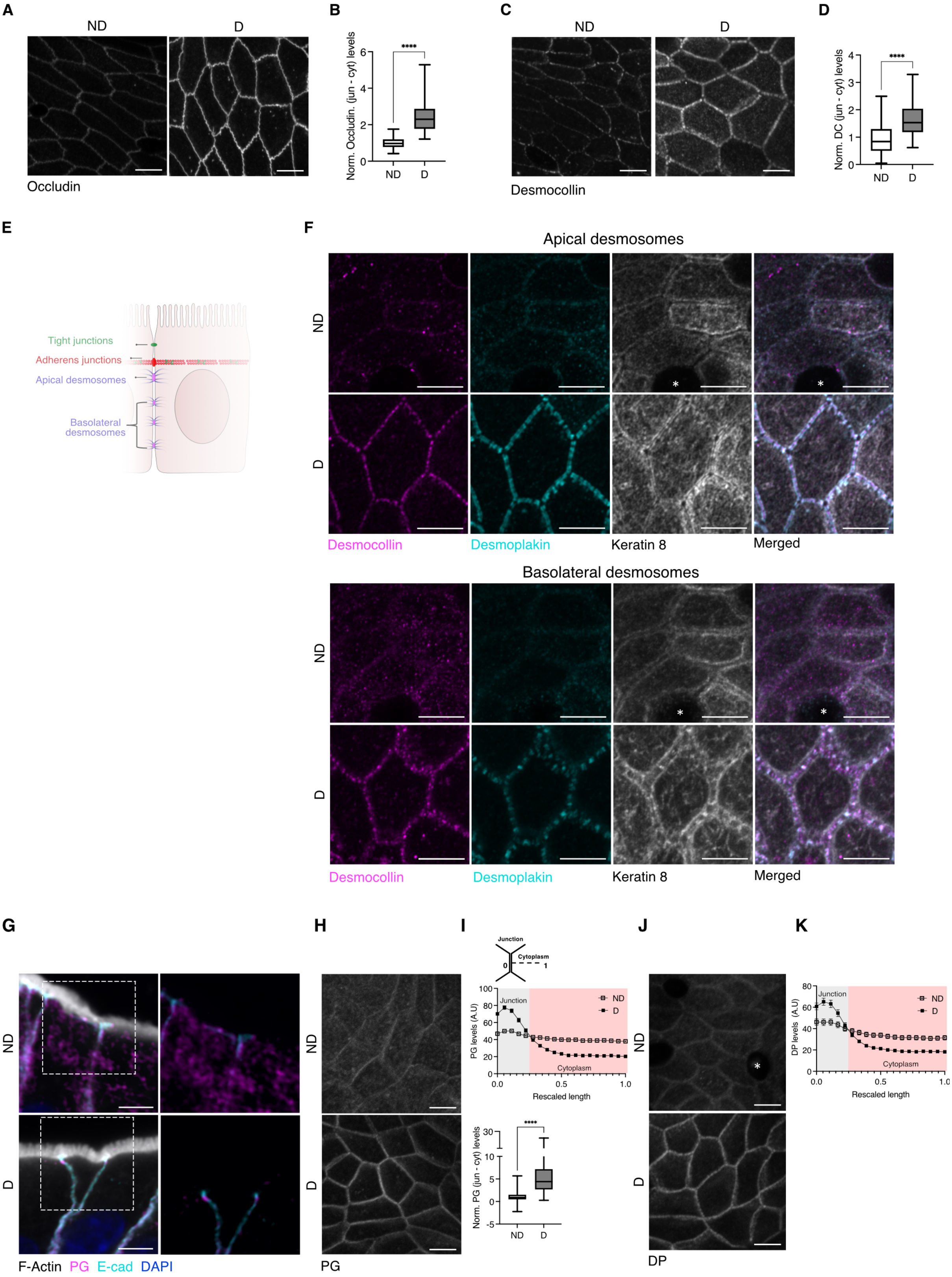


**Fig. S1**. **(A)** Representative *en face* images of colonic epithelium, stained for occludin in ND and D regions. Maximum Z-projections, 1-3 $\mu$m. Scale bar: 5 $\mu$m. **(B)** Box plots showing junctional intensity for occludin (n=48-51 junctions/condition) in ND and D regions, N=3 experiments **(C)** Representative *en face* images of colonic epithelium, stained for desmocollin in ND and D regions. Maximum Z-projections, 1-3 $\mu$m. Scale bar: 5 $\mu$m. **(D)** Box plots showing junctional intensity for desmocollin (n=53-73

junctions/condition) in ND and D regions, N=3 experiments. **(E)** Scheme showing adhesive junctional complexes (TJs, AJs and Ds). The apical desmosomes are located immediately below the E-cad (AJ) level. Basolateral desmosomes are located below the most apical desmosomes. **(F)** Representative *en face* images of colonic epithelium stained for DC (magenta), DP (cyan) and K8 (grey) showing apical desmosomes (top); maximum Z-projection (1-3 $\mu$m range) and basolateral desmosomes (bottom); maximum Z-projection (4-6 $\mu$m range). Scale bars: 5 $\mu$m. **(G)** Representative transverse images of colonic epithelium, stained for F-actin (grey), PG (magenta), E-cad (cyan) and DAPI (blue) in both ND and D regions. Maximum Z-projections (2-4 $\mu$m range). Dashed rectangle indicates region shown at higher magnification. Scale bars: 5 $\mu$m. **(H)** Representative *en face* images of colonic epithelium, stained for PG in ND and D regions. Maximum Z-projections (1-3 $\mu$m range). Scale bar: 5 $\mu$m. **(I)** Top: Scheme showing plot profile measurement, briefly a line of fixed length is drawn from junction to cytoplasm and intensity is measured along the line. Middle: Plot profile for PG in ND and D regions of colonic epithelium. Bottom: Box plot showing junctional intensity for PG in ND and D regions (n=28-54 junctions/condition, N=7 independent experiments). **(J)** Representative *en face* images of colonic epithelium, stained for DP in ND and D regions. Maximum Z-projections, 1-3 $\mu$m. Scale bar: 5 $\mu$m. **(K)** Plot profile for DP in ND and D regions of colonic epithelium. A line of fixed length is drawn from junction to cytoplasm and intensity is measured along the line. Statistical analyses were performed using the Mann–Whitney U test (B), (D) and (I). Significance is denoted as: ****$p < 0.0001$, ***$p < 0.0005$, ** $p < 0.005$, ns: non-significant. ND: Non-distended, D: Distended, R: Relaxed. *: goblet cell.
or Kruskal-Wallis test followed by Dunn's multiple comparison (J) and (M).

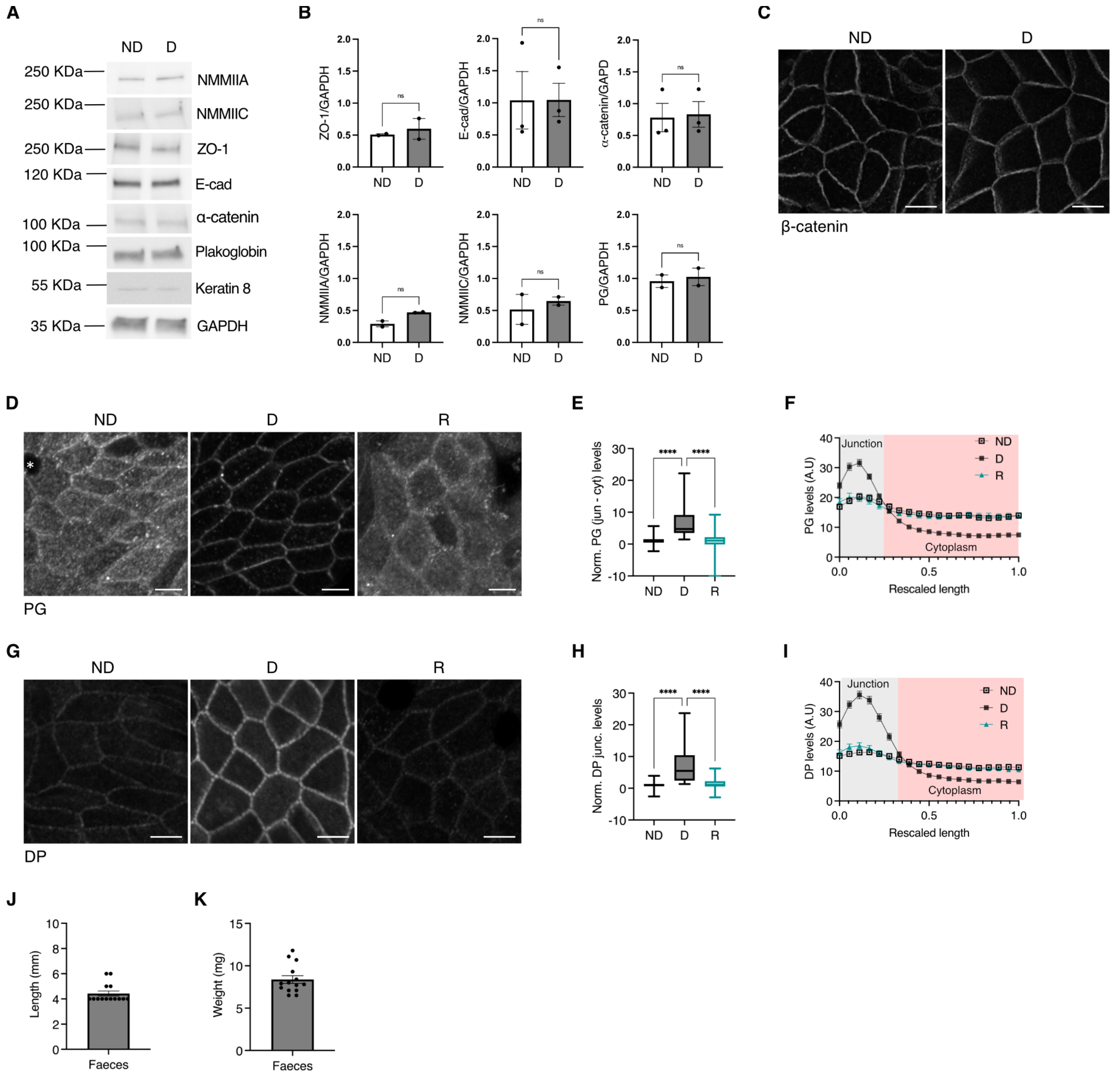

**Fig. S2**. **(A-B)** Representative (A) western blot images and (B) quantification for NMMIIA, NMMIIC, ZO-1, E-cad, α-catenin, PG, K8 and GAPDH from protein lysates from ND or D regions of the colonic epithelium. GAPDH was used as a loading control. N=2 independent experiments. **(C)** Representative *en face* images of colonic epithelium stained for β-catenin, showing ND, D regions. Maximum Z-projections (1-3 $\mu$m range). Scale bar: 5 $\mu$m. **(D)** Representative *en face* images of colonic epithelium stained for PG, showing ND, D and R regions. Maximum Z-projections, 1-3 $\mu$m. Scale bar: 5 $\mu$m. **(E-F)** Box plot showing: (E) junctional PG intensity in ND, D and R regions (n=44-54 junctions/condition, N=3 independent experiments). (F) Plot profile for PG in ND, D and R regions of colonic epithelium. **(G)** Representative *en face* images of colonic epithelium stained for DP, showing ND, D and R regions. Maximum Z-projections (1-3 $\mu$m range). Scale bar: 5 $\mu$m. **(H-I)** Box plot showing: (H) junctional DP intensity in ND, D and R regions (n=48-63 junctions/condition, N=3 independent experiments). (I) Plot profile for DP in ND, D and R regions of colonic epithelium. **(J-K)** Bar plots showing length of the faeces pellet(C) and weight of the faeces pellet (D). (n=15 faeces pellets; N=5 independent experiments). Statistical analyses were performed using the Mann–Whitney U test (B) or Kruskal-Wallis test followed by Dunn's multiple comparison (E) and (H). Significance is denoted as: ****p < 0.0001, ***p < 0.0005, ** p < 0.005, ns: non-significant. ND: Non-distended, D: Distended, R: Relaxed. *: goblet cell.

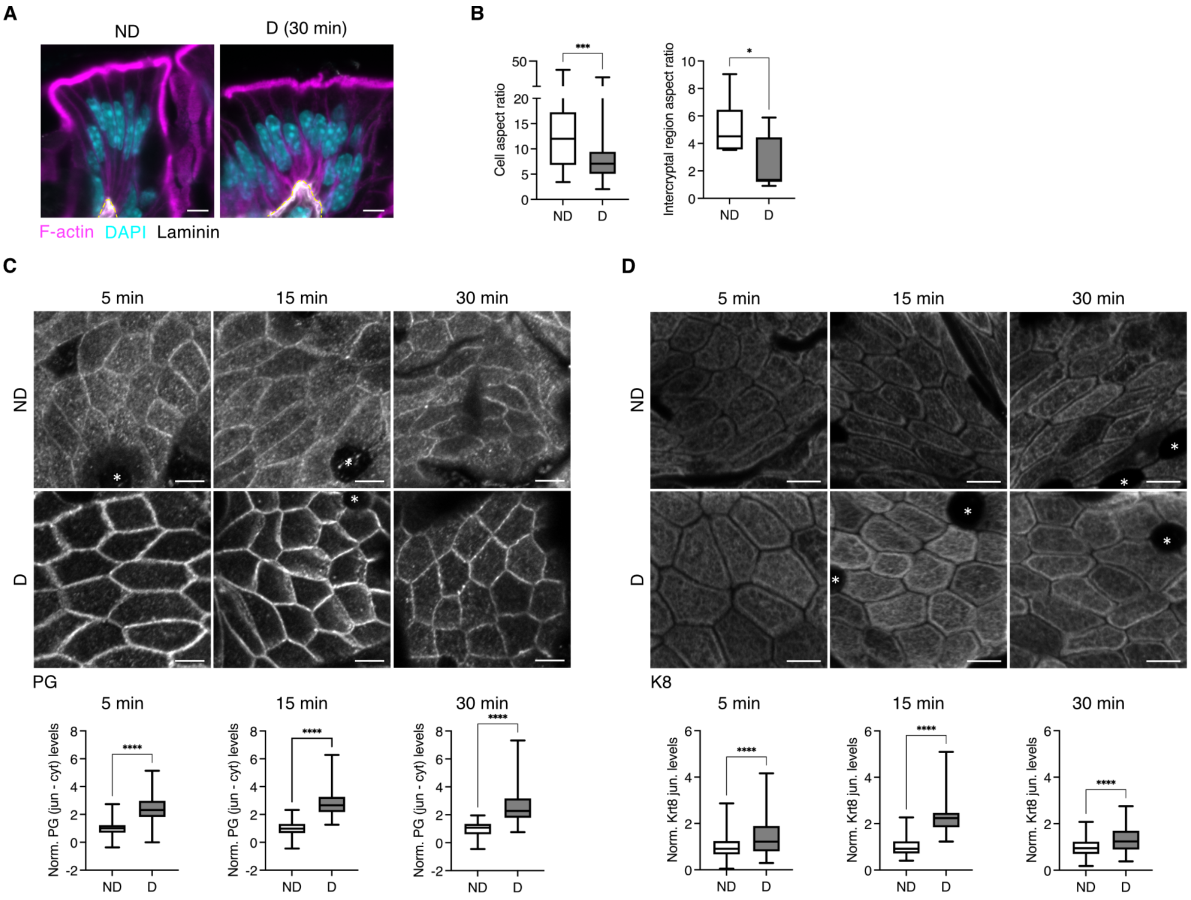


**Fig. S3 (A)** Representative transverse images of plateau regions in the colonic epithelium, stained for F-actin (magenta), DAPI (cyan) and laminin (grey) in both non-distended (ND) and catheter-mediated distended (D) regions. Maximum Z-projections (2-4 $\mu$m range). Scale bar: 5 $\mu$m. **(B)** Box plots showing: cell aspect ratio (left; n=15-24 cells/condition) and intercryptal aspect ratio (right; n=3-4 intercryptal regions/condition) in ND and D plateau regions. N=2 independent experiments. **(C)** Top: Representative *en face* images of colonic epithelium, stained for PG in ND and D regions after 5,15 and 30 min of distension. Maximum Z-projections (1-3 $\mu$m range). Scale bar: 5 $\mu$m. Bottom (left to right): Box plots showing junctional PG intensity in ND and D regions after 5, 15 and 30 min of distension. (n=42-57 junctions/condition, N=3-5 independent experiments). **(D)** Top: Representative *en face* images of colonic epithelium, stained for K8 in ND and D regions after 5,15 and 30 min of distension. Maximum Z-projection (1-3 $\mu$m range). Scale bar: 5 $\mu$m. Bottom (left to right): Box plots showing junctional K8 intensity in ND and D regions after 5, 15 and 30 min of distension. (n=28-84 junctions/condition, N=4-3 independent experiments). Statistical analyses were performed using the Mann–Whitney U test or welch-t-test (B, intercryptal aspect ratio). Significance is denoted as: ****$p < 0.0001$, ***$p < 0.0005$, ** $p < 0.005$, ns: non-significant. ND: Non-distended, D: Catheter-mediated distended. *: goblet cell.

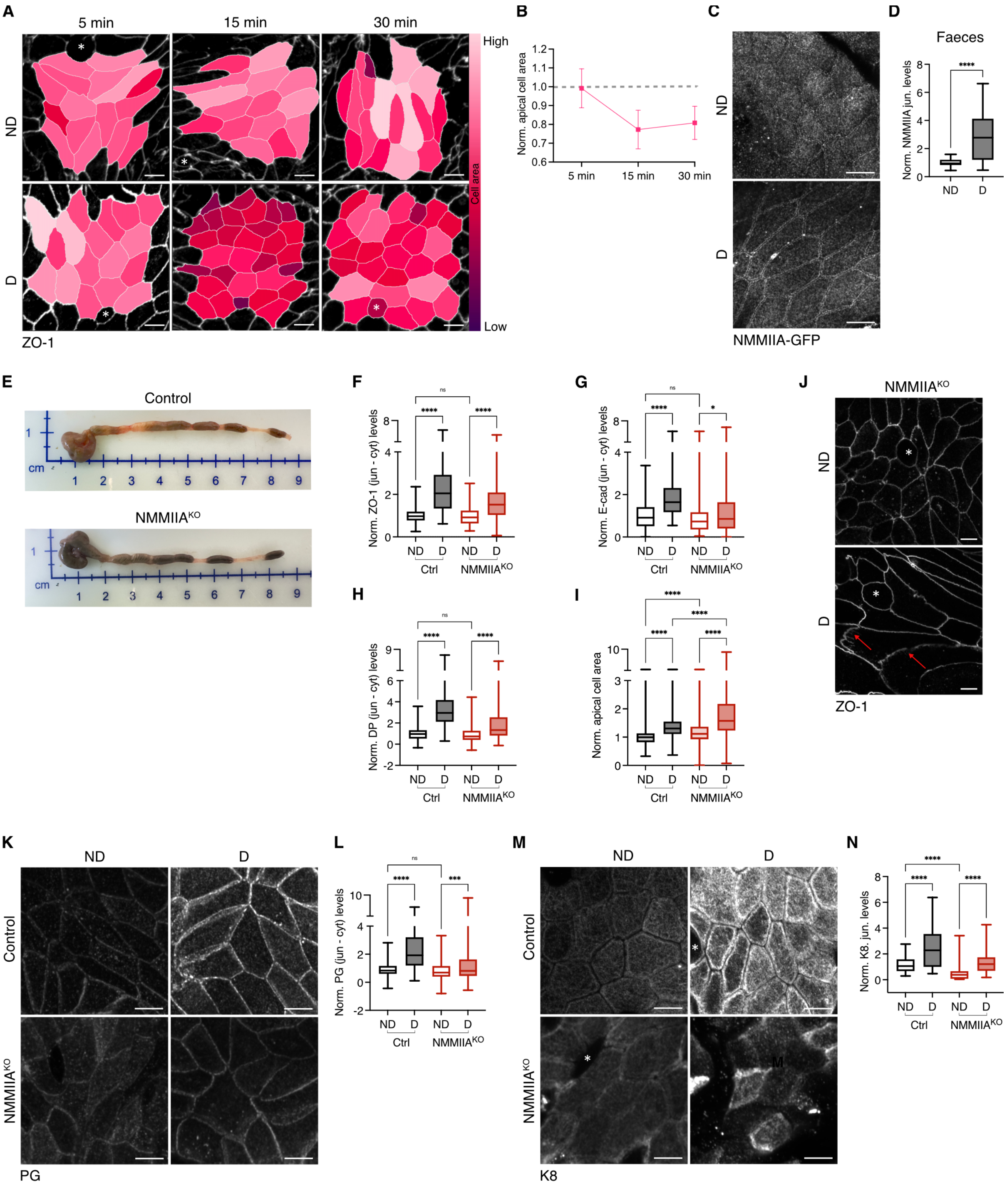


**Fig. S4. (A)** Representative images, segmented and colour-coded based on apical cell area in ND and D regions after 5,15 and 30 min of distension. Maximum Z-projections (1-3 $\mu$m range). Scale bar: 5 $\mu$m. **(B)** Line plot showing mean fold change of apical cell area after 5, 15 and 30 min of distension. Dotted line represents the normalised average for ND. **(C)** Representative *en face* images of colonic epithelium showing NMMIIA-GFP in ND and faeces-D regions. Maximum Z-projection (1-3 $\mu$m range). Scale bar: 5 $\mu$m. **(D)** Box plot showing junctional NMMIIA-GFP intensity in ND and faeces-D regions (n=18-32 junctions/condition, N=4 independent experiments). **(E)** Representative images of mouse colon, aligned from the proximal (caecum) to the distal (rectum) end, in control and NMMIIA$^{KO}$ mice. **(F-H)** Box plots showing junctional intensity levels in ND and D regions for ZO-1 (H), E-cad (I), and DP (J) in control and NMMIIA$^{KO}$ (n=38-84 junctions/condition, N=4-5 independent experiments) **(I)** Box plot showing apical cell area in control and NMMIIA$^{KO}$ mice normalized to control ND values (n=42-544 cells/condition, N=3 independent experiments) **(J)** Representative images of NMMIIA$^{KO}$ colonic epithelium stained for ZO-1 in ND and faeces-D regions. Red arrows represent discontinuous ZO-1 staining. Maximum Z-projection (1-3 $\mu$m range). Scale bar: 5 $\mu$m. **(K)** Representative images of colonic epithelium stained for PG in control and NMMIIA$^{KO}$ from ND and faeces-D regions. Maximum Z-projection (1-3 $\mu$m range). Scale bar: 5 $\mu$m. **(L)** Box plots showing junctional intensity levels for PG in control and NMMIIA$^{KO}$, normalized to ND control values (n=60-77 junctions/condition, N=5 independent experiments). **(M)** Representative images of colonic epithelium stained for K8 in control and NMMIIA$^{KO}$ from ND and faeces-D regions. Maximum Z-projection (1-3 $\mu$m range). Scale bar: 5 $\mu$m. **(N)** Box plots showing junctional intensity levels for K8 in control and NMMIIA$^{KO}$, normalized to ND control values (n=45-54 junctions/condition, N=3 independent experiments). Statistical analyses were performed using the Mann–Whitney U test (D) or Kruskal-Wallis test followed by Dunn's multiple comparison (F-I), (L) and (N). Significance is denoted as: ****p < 0.0001, ***p < 0.0005, ** p < 0.005, ns: non-significant. ND: Non-distended, D: Distended. *: Indicates goblet cell.

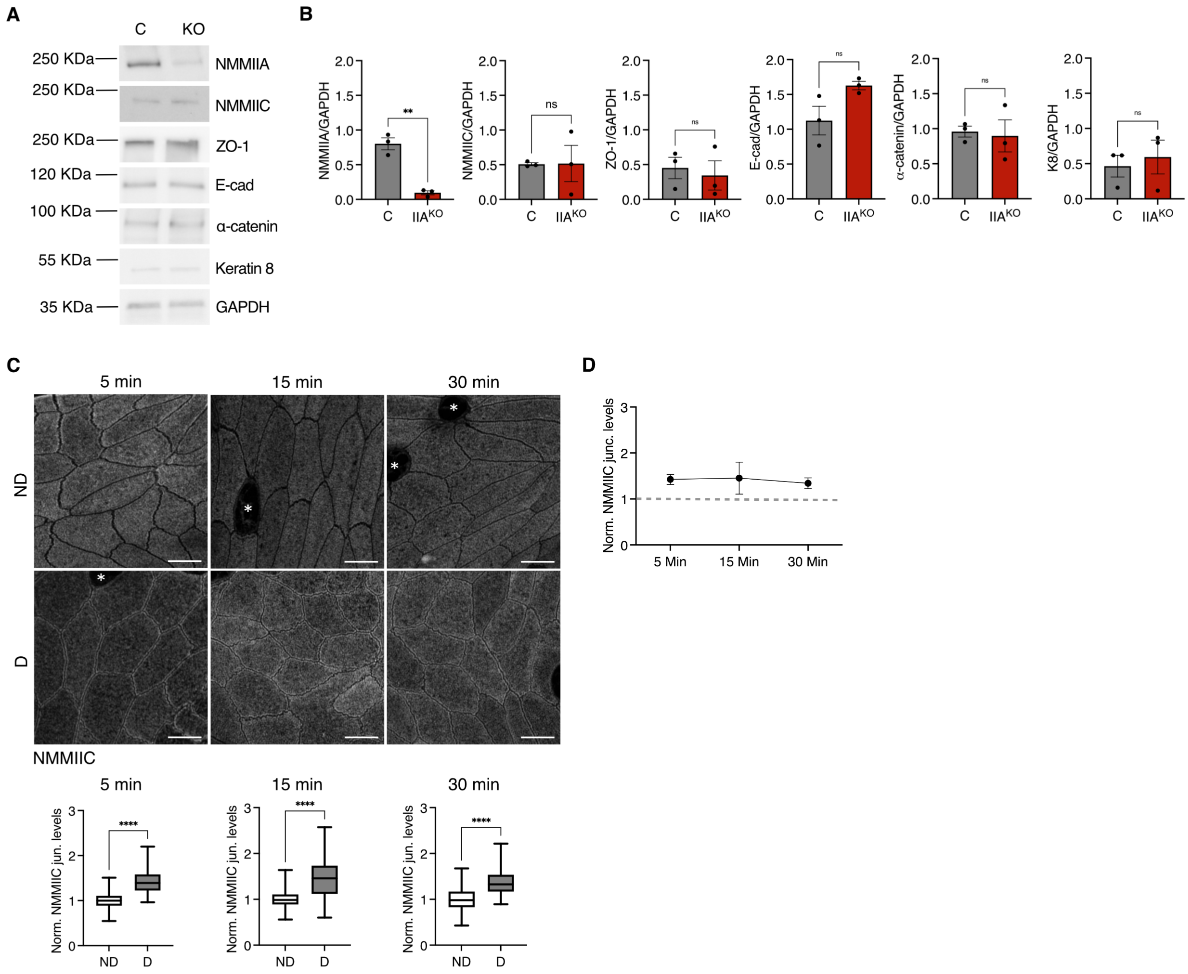


**Fig. S5. (A)** Representative western blot images and **(B)** quantification for NMMIIA, NMMIIC, ZO-1, E-cad, α-catenin, K8 and GAPDH from protein lysates isolated from ND and faeces-D regions of control (C) and NMMIIA$^{KO}$ (KO) mice. GAPDH was used as a loading control (N=3 independent experiments). **(C)** Top: Representative *en face* images of colonic epithelium, stained for NMMIIC in non-distended (ND) and catheter-distended (D) regions after 5, 15 and 30 min of distension. Maximum Z-projection (1-3 $\mu$m range). Bottom (left to right): Box plots showing junctional NMMIIC intensity in ND and D regions after 5, 15 and 30 min of distension (n=48-64 junctions/condition, N=2-3 independent experiments). **(D)** Line plot showing mean fold change in NMMIIC intensity after 5, 15 and 30 min of distension. Dotted line represents the normalised average for ND. Statistical analyses were performed using the Welsch-t test (B) or Mann-Whitney (C). Significance is denoted as: ****p < 0.0001, ***p < 0.0005, ** p < 0.005, ns: non-significant. ND: Non-distended, D: Distended. *: goblet cell. Scale bar: 5 $\mu$m.

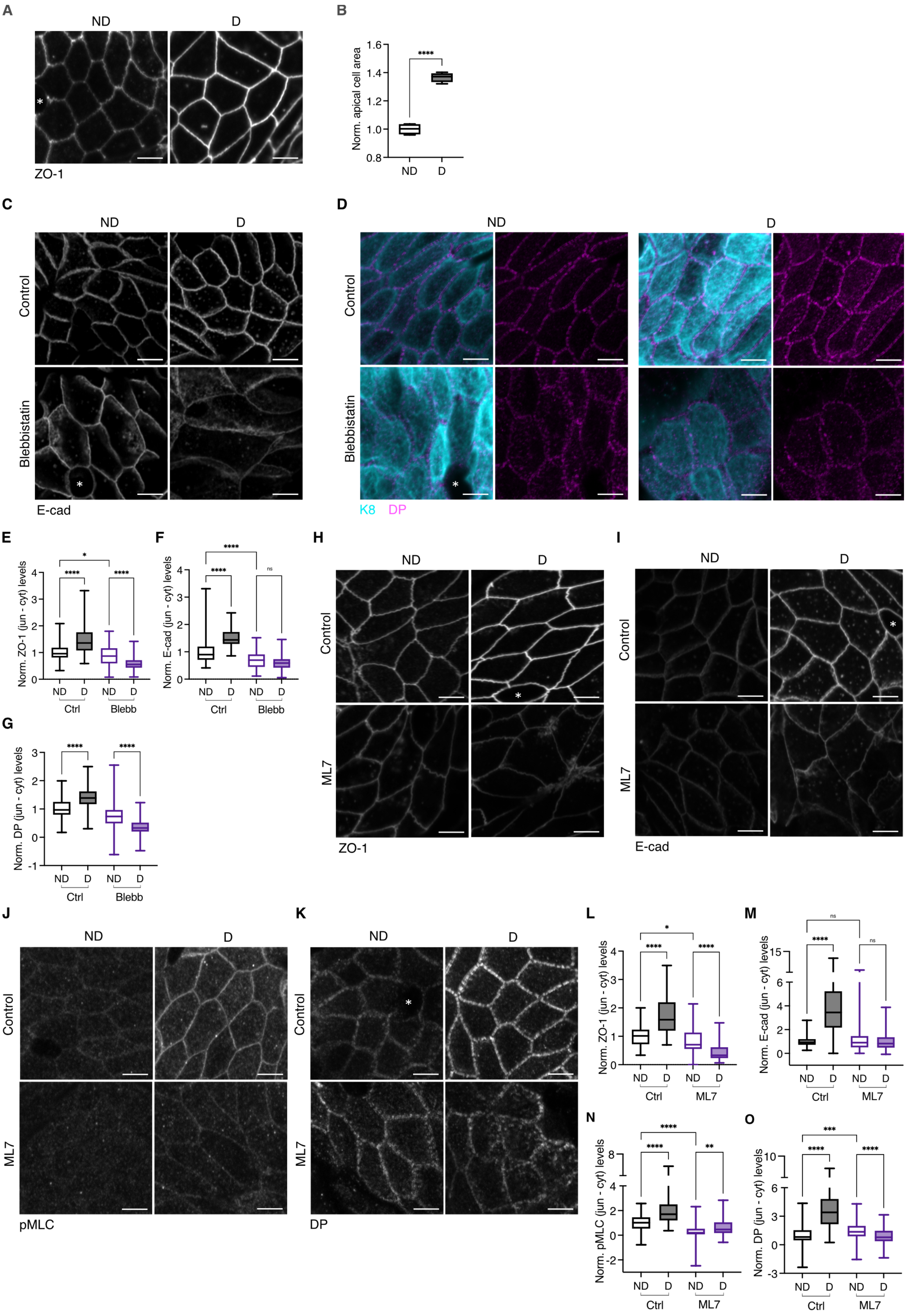


**Fig. S6. (A)** Representative images showing colonic explants stained for ZO-1 in ND and D after 30 min of distension. Maximum Z-projection (1-3 $\mu$m range). Scale bar: 5 $\mu$m. **(B)** Box plot showing apical cell area in D-colon explants, normalized to ND regions (n=22-37 cells/condition, N=4 independent experiments). **(C-D)** Representative images showing colonic explants stained for E-cad (C), and K8 (cyan) and DP (magenta) (D) in ND and D explants after 1 h pre-treatment with blebbistatin or vehicle control (DMSO) followed by 30 min distension. Maximum Z-projection (1-3 $\mu$m range). Scale bar: 5 $\mu$m. **(E-G)** Box plots showing junctional intensities for ZO-1 (E), E-cad (F) and DP (G) in ND and D explants after pre-treatment with blebbistatin or Vehicle control (DMSO) and distension (n=30-79 junctions/condition, N=2-3 independent experiments). **(H-K)** Representative images showing colonic explants stained for ZO-1 (H), E-cad (I), pMLC (S19) (J) and DP (K) in ND and D explants after 1 h pre-treatment with ML-7 or vehicle control (DMSO) followed by 30 min distension. Maximum Z-projection (1-3 $\mu$m range). Scale bar: 5 $\mu$m. **(L-O)** Box plots showing junctional intensity for ZO-1 (L), E-cad (M), pMLC (N) and DP (O) in ND and D explants after pre-treatment with ML-7 or vehicle control (DMSO) and distension (n=39-74 junctions/condition, N=3-5 independent experiments). Statistical analyses were performed using the Welch t-test (B) or Kruskal-Wallis test followed by Dunn's multiple comparison (E-G) and (L-O). Significance is denoted as: ****$p < 0.0001$, ***$p < 0.0005$, ** $p < 0.005$, * $p < 0.05$, ns: non-significant. ND: Non-distended, D: Distended. *: goblet cell.

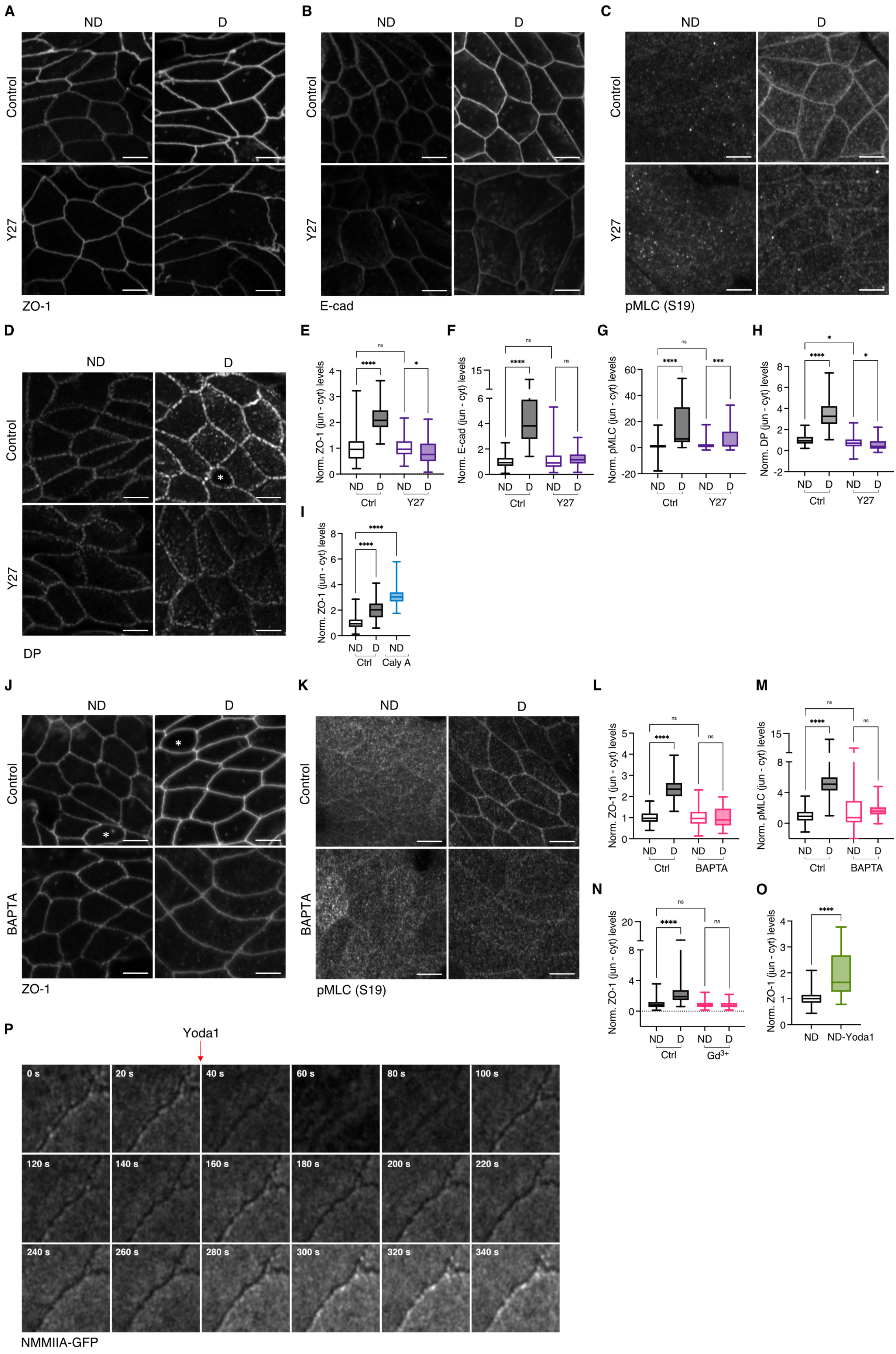

A
ND
D
Control
Y27
ZO-1
B
E-cad
C
pMLC (S19)
D
DP
E
Norm. ZO-1 (jun - cyt) levels
ns
****
*
ND D ND D
Ctrl Y27
F
Norm. E-cad (jun - cyt) levels
G
Norm. pMLC (jun - cyt) levels
***
H
Norm. DP (jun - cyt) levels
I
Caly A
J
BAPTA
K
L
M
N
Gd3+
O
ND-Yoda1
P
Yoda1
0 s
20 s
40 s
60 s
80 s
100 s
120 s
140 s
160 s
180 s
200 s
220 s
240 s
260 s
280 s
300 s
320 s
340 s
NMMIIA-GFP

**Fig. S7**. **(A-D)** Representative images showing colonic explants stained for ZO-1 (A), E-cad (B), pMLC (S19) (C) and DP (D) in ND and D regions after 1 h pre-treatment with Y27632 or vehicle control (water), followed by 30 min distension. Maximum apical Z-projection (1-3 $\mu$m range). Scale bar: 5 $\mu$m. **(E-H)** Box plots showing junctional intensity for ZO-1(E), (F) E-cad, pMLC (S19) (G) and DP (H) in ND and D explants after pre-treatment with Y27632 (Y27) or vehicle control (water) and distension (n=31-54 junctions/condition, N=3-4 experiments). **(I)** Box plot showing junctional ZO-1 intensity in Calyculin A-treated ND explants and vehicle-treated ND and D explants (n=53-59 junctions/condition, N=3 independent experiments). **(J-K)** Representative image showing colonic explants stained for ZO-1 (I) and pMLC (S19) (J) in ND and D after treatment with BAPTA or vehicle control (water), followed by 30 min distension. Maximum Z-projection (1-3 $\mu$m range). Scale bar: 5 $\mu$m. **(L-M)** Box plots showing junctional intensity for ZO-1 (K) and pMLC (S19) (L) in ND and D explants after treatment with BAPTA or vehicle control (water) and distension (n=47-54 junctions/condition, N=2 independent experiments). **(N)** Box plot showing junctional ZO-1 intensity in ND and D explants after pre-treatment with $Gd^{3+}$ or vehicle control (water) and distension (n=58-61 junctions/condition, N=5 independent experiments). **(O)** Box plot showing junctional ZO-1 intensity in ND explants after treatment with Yoda1 and vehicle control (DMSO) (n=55-66 junctions/condition, N=2 independent experiments). **(P)** Representative montage showing junctional recruitment of NMMIIA after addition of Piezo-1 activator, Yoda1. Statistical analyses were performed using the Mann-Whitney U test (O) or Kruskal-Wallis test followed by Dunn's multiple comparison (E-H) and (K-N). Significance is denoted as: ****$p < 0.0001$, ***$p < 0.0005$, ** $p < 0.005$, ns: non-significant. ND: Non-distended, D: Distended.

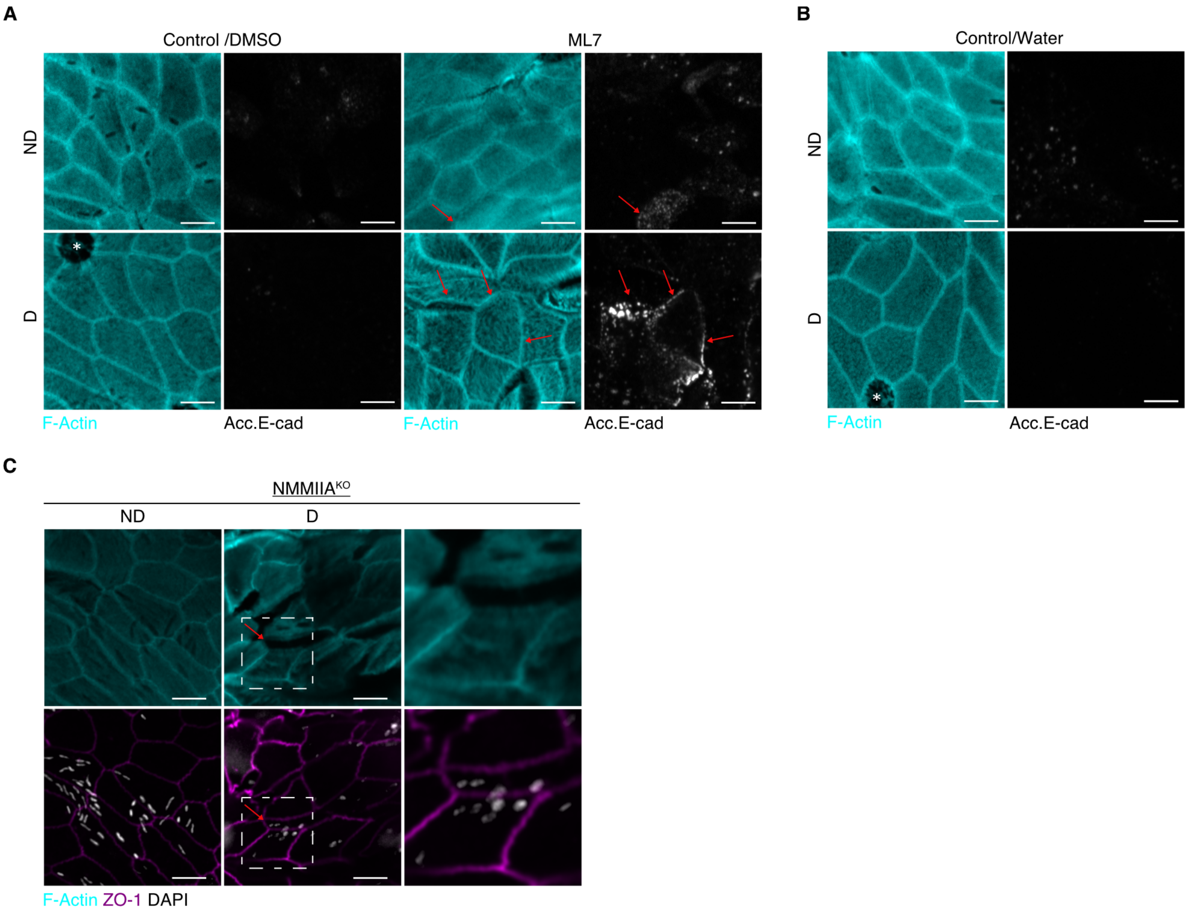


**Fig. S8. (A)** Representative images showing colonic explants stained for F-actin (cyan) and acc. E-cad (grey) in ND and D after 1 h pre-treatment with ML7 or vehicle (DMSO)followed by 30 min distension. Red arrows indicate breached junctions. Maximum apical Z-projection (1-3 $\mu$m range). *: goblet cell. Scale bar 5 $\mu$m. **(B)** Representative image showing colonic explants stained for F-actin (cyan) and acc. E-cad (grey) in ND and D after 1 h pre-treatment with vehicle (water) followed by 30 min distension. Maximum Z-projection (1-3 $\mu$m range). *: goblet cell. Scale bar: 5 $\mu$m. **(C)** Representative image of colonic epithelium stained for F-actin (cyan), ZO-1 (magenta) and DAPI (grey) in NMMIIA$^{KO}$ mice from ND and faeces-D regions. Red arrows indicate translocation of bacteria inside the fractured regions. Dashed rectangle indicates region shown at higher magnification. Maximum Z-projection (1-3 $\mu$m range). Scale bar: 5 $\mu$m.
